# HybriSeq: Probe-based Device-free Single-cell RNA Profiling

**DOI:** 10.1101/2023.09.27.559406

**Authors:** Daniel Foyt, David Brown, Shuqin Zhou, Brittany Moser, Qin Zhu, Zev J Gartner, Bo Huang

## Abstract

We have developed the HybriSeq method for single-cell RNA profiling, which utilizes *in situ* hybridization of multiple probes for targeted transcripts, followed by split-pool barcoding and sequencing analysis of the probes. We have shown that HybriSeq can achieve high sensitivity for RNA detection with multiple probes and profile entire transcripts without an end bias. The utility of HybriSeq is demonstrated in characterizing cell-to-cell heterogeneities of a panel of 196 genes in peripheral blood mononuclear cells and the detection of missed annotations of transcripts.

## Introduction

With its ability to profile individual transcriptomes of many cells, single-cell RNA sequencing (scRNAseq) has proven to be an invaluable tool in understanding cell-to-cell heterogeneity and gene regulatory networks in complex systems (1, 2, 3). Most scRNAseq methods capture polyadenylated RNA and then use reverse transcription to convert it into double-stranded DNA that is compatible with sequencing reactions (4). Although this approach can analyze mRNAs in an unbiased way, the typical detection efficiencies for individual RNA transcripts range between 5-45% (5, 6, 7), largely caused by the inefficiency of the template-switching reaction during reverse transcription, inefficient priming, and RNA degradation. These inefficiencies are particularly deleterious for detection of low copy number RNA and lead to drop out or noisy measurements making classification of subtle phenotypes difficult with few cells (8).

In contrast to the low detection efficiency in scRNAseq, single-molecule fluorescence *in situ* hybridization (smFISH) regularly achieves a detection efficiency close to 100% by utilizing multiple probes to probe the target RNA directly (9). Taking this concept, single-cell RNA profiling can also be achieved by sequencing multiple *in situ* hybridization probes for one given transcript to decrease the likelihood of a molecule going undetected and increase the measurement confidence. Indeed, several probe-based single-cell RNA profiling methods have been developed recently, such as HyPR-Seq (10), ProBac-seq (11), and 10X Genomics Chromium Flex protocol (12). Due to their probe-based nature, these methods are inherently targeted, allowing for efficient utilization of sequencing reads, and they are not limited to profiling polyadenylated RNA like many scRNAseq methods. On the other hand, they each have their unique limitations. For instance, their probe chemistry either requires complex oligo hybridization and ligation steps, leading to low probe detection efficiency and high background, or simply relies only on hybridization-based specificity, leading to low specificity. Additionally, all of them use microfluidic partitioning of single cells, which can limit the number of cells profiled and require costly instrumentation. In contrast, highly scalable methods such as SPLiT-Seq (13) and Sci-Plex (14) can sequence millions of cells by utilizing combinatorial indexing. Combining a probe-based approach with combinatorial indexing can thus enable single-cell RNA profiling with both high sensitivity/efficiency and high throughput.

In pursuit of this goal, we developed **Hybri**dization of probes to RNA targets followed by **Seq**uencing (**HybriSeq**). This method involves *in situ* hybridization of multiple split single-strand DNA (ssDNA) probes to one or many target RNAs in fixed and permeabilized cells (Fig. 1a), ligating these split probes hybridized to the RNA to ensure specificity (Fig. 1b), ligating a unique cell barcode to the hybridized probes via two rounds of split-pool barcoding followed by an indexed polymerase chain reaction (PCR) (Fig. 1c), and sequencing the ligated probe-barcodes. This method can sensitively detect transcripts in a targeted fashion without the need for microfluidics. We demonstrate the utility of this method by profiling the cell-cell heterogeneity in peripheral blood mononuclear cells (PBMC).

**Fig. 1.**
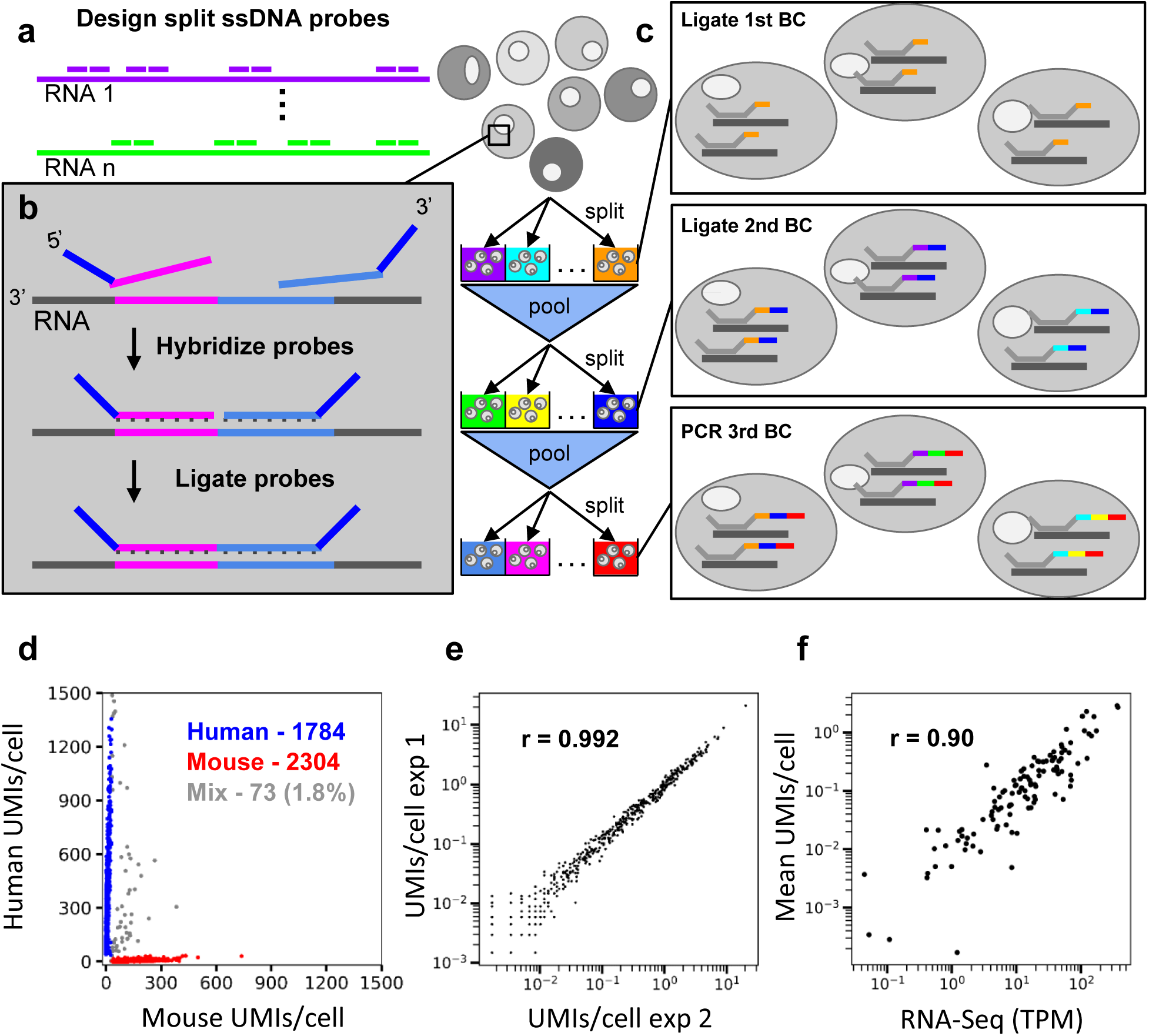
Schematic and Validation of HybriSeq. **a.** Multiple split probes are designed per transcript of interest. **b.** Hybridization and ligation of split probes. **c.** Labeling probes with unique cell barcodes via the split-and-pool method. Rounds 1 and 2 barcodes are ligated, and round 3 barcodes are added via PCR. **d.** Sequencing results of 1:1 mixed HEK293 cells and Do-11-10 cells. 1.8% of barcodes contain probes targeting human and mouse transcripts. 98.2% of barcodes contain probes targeting only human or only mouse transcripts. **e.** Scatter plot of average HybriSeq UMIs per cell in two independent biological replicates. Each dot represents a single probe. Pearson Correlation Coefficient (r = 0.992, p < 0.0001) **f.** Scatter plot of gene level average HybriSeq UMIs per cell and expression of the same gene obtained via bulk RNA-Seq. Each dot represents a gene level measurement. Pearson Correlation Coefficient (r = 0.90, p < 0.0001)

## Results

### Development and Validation of HybriSeq

To establish a method for efficient hybridization and recovery of ssDNA probes to target RNAs with low nonspecific binding, we performed *in situ* hybridization in fixed and permeabilized human embryonic kidney 293 (HEK293) cells in suspension and quantified the efficiency and specificity of probe recovery by sequencing. We found that ssDNA probes have non-negligible nonspecific binding to the cells, which can contribute to background signal (Supplementary Fig. 1b). We tried to improve the specificity by selectively releasing hybridized probes using RNaes H digestion of the cell’s RNA (Supplementary Fig. 1c), but the signal/background ratio was still low even after optimizing hybridization conditions (Supplementary Figs. 1d-e) (see Supplementary Note 1). Therefore, we adopted a method similar to LISH (15), MLPA (16), and ClampFISH (17), splitting the probe into two parts and ligating hybridized pairs using SplintR ligase that acts on DNA-RNA hybrids (Figs. 1a-b). Bulk-level quantitative polymerase chain reaction (qPCR) measurements of ligated probes in cells showed that with ligation (Supplementary Fig. 2a), it is possible to saturate the probe signal from a moderately expression transcript (Supplementary Figs. 2b-c) and achieve a specific signal > 1000 times higher than nontargeting probes (Supplementary Fig. 2b) (see Supplementary Note 1).

To enable single-cell analysis, we adapted the split-pooling method (13) to uniquely label the probes in individual cells with cell-specific barcodes (see Supplementary Note 2). In 96-well plates, hybridized and ligated probes are labeled with well-specific barcodes via ligation on the 3’ end in two rounds of split and pool procedures followed by a third round of barcoding by PCR with well specific primers (Fig. 1c). Depending on the path a cell takes through this procedure, all the probes in that cell will have one of 884,736 possible unique cell barcodes (CBC), of which < 5% are utilized to avoid excessive CBC collision, a situation in which multiple cells take identical paths resulting in a CBC associated with multiple cell’s transcriptomes. Different from previous split-pooling methods, our main challenge is the mixing of barcodes between cells, which can arise from inefficient ligation before pooling, excess unhybridized barcode oligos in subsequent ligation reactions, which can hybridize and be ligated incorrectly, and priming of incompletely ligated species. We screened a variety of barcode ligation and washing/quenching conditions in bulk with qPCR. We found that long ligation times and high barcode oligo concentrations are needed for efficient barcode ligation (Supplementary Fig. 3b). Additionally, we found that quenching barcode oligos as opposed to blocking linker strands resulted in less barcode hopping and that washing away excess barcodes after each ligation step led to significantly less barcode hopping (Supplementary Figs. 3c-f).

To investigate the performance and single-cell purity of HybriSeq, we designed a set of probes (2-4 probes per transcript) targeting 95 human cell cycle-associated transcripts and 124 mouse cell cycle-associated transcripts. (Supplementary Table 1). We used this probe set to perform a cell mixing experiment in which we mixed HEK293 cells and mouse helper T cells (Do-11-10) in equal proportions before performing HybriSeq. We observed that 1.8% of CBC contained multiple probes from both mouse and human transcripts, suggesting a doublet rate of 3.6% (Fig. 1d) (from a 50/50 mix of cells, half of the doublets will arise from two cells with the same CBC). Unique molecular identifier (UMI) counts per probe per cell in a similar but separate experiment were highly correlated between biological replicates with a Pearson correlation coefficient >0.99 (Fig. 1e). A median of 97.3% of reads for each CBC were specific to either mouse or human probes. These data suggest HybriSeq libraries have a high level of single-cell purity and reproducibility. This multiple rate is higher than the expected multiple rate of 2.45%, which is most likely due to cell clumping. While nonzero, this value is lower than most droplet-based approaches.

The specificity of HybriSeq arises from both the specific hybridization of ssDNA to transcripts and from the ligation of two adjacent probes hybridized. To evaluate the specificity of HybriSeq, we examined reads in the library that contained left-probe and right-probe targeting regions not predicted to be adjacent to each other. We compared the amount of these nonspecific ligation events to the specific and correctly ligated events. Nonspecific ligation events accounted for an average of 0.20% of UMIs per cell in cells with a median of 150 UMIs per cell (Supplementary Fig. 4a-b). This result suggests that HybriSeq is highly specific.

To demonstrate the quantitative accuracy of HybriSeq, we picked one of the two cell lines and probe sets from the cell mixing experiment (Do-11-10 cell line and mouse transcript probes) and performed both HybriSeq and standard RNA-Seq analysis. The resulting bulk expression values correlate well with matched sample bulk RNA-Seq data (*r* = 0.90) (Fig. 1f). To determine the effect on measurement precision if fewer probes per transcript were used, we subsampled the number of probes used to calculate a Pearson correlation coefficient. As expected, the use of 3 probes per transcript improved correlation, while less accurate results were seen when 1-2 probes were sampled (Supplementary Fig. 4c).

### Variability of Probe Detection Efficiency

To characterize the variability of detection efficiency for probes designed using our method, we constructed a set of probes completely tiling 6 transcripts (tiling probes) with expression levels ranging from 15 to 165 Transcripts per million (TPM) in HEK293 cells (Supplementary Fig. 5, Supplementary Table 2). These transcripts are expected to only have one isoform expressed that does not have expected variation during the cell cycle, which is the main source of heterogeneity in a monoculture cell culture system. Our transcript tiling results from profiling 394 cells reveal that the vast majority of probes targeting the same transcript have low variabilities in the average detected number of UMIs per cell (Fig. 2a-f). For example, SCAF8, ARL5B, and MARVELD1 showed a relatively uniform probe occupancy throughout the length of the transcript with few outliers (Figs. 2c-e). Surprisingly, for the two transcripts with high expression, we observed that continuous stretches of probes are underrepresented or hardly represented at all, despite the otherwise highly uniform probe representation (Fig. 2a-b). For EIF2S2 (ENSG00000125977), the 3’ half of the transcript has very few UMIs associated with it (Fig. 2a). While not as extensive, this depletion is also seen in GHITM (ENSG00000165678) (Fig. 2b).

**Fig. 2.**
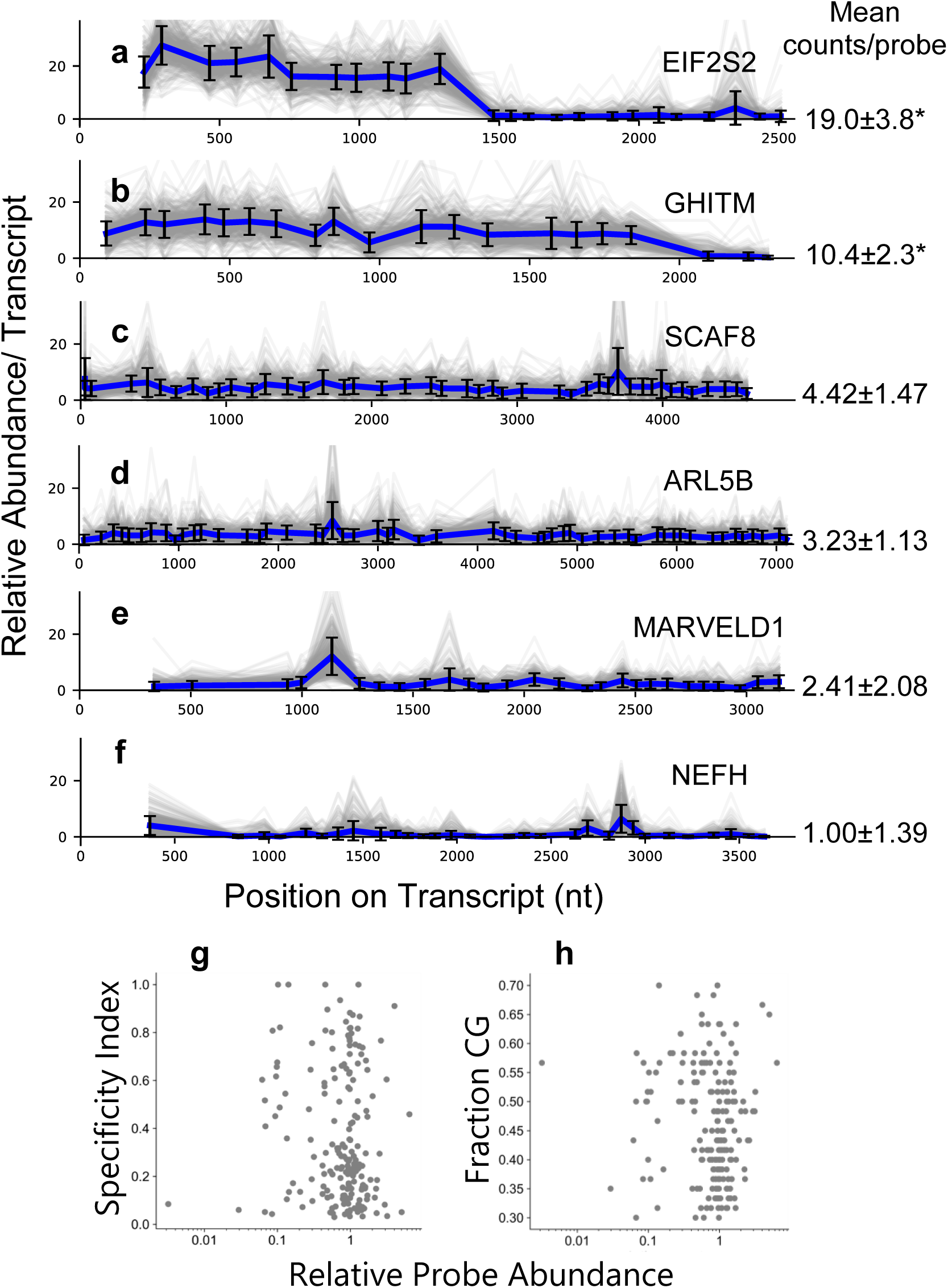
Intra-transcript Probing Variation. a-f. Relative probe counts for probes targeting the specific transcript with standard deviation across all 394 cells plotted for each probe. Gray lines are traces for individual cells, and the blue line is the average for each probe across all cells. The means of average counts for all probes reported to the right with standard deviations. (*) For EIF2S2, only the first 11 out of 24 probes were considered in calculating the mean and standard deviation. For GHITM only the first 17 out of 20 probes were considered in calculating the mean and standard deviation. **g.** Average relative probe abundance normalized for each transcript for individual probes plotted against calculated probe specificity index (see HybriSeq Split probe design in methods) **h.** Average relative probe abundance normalized for each transcript for individual probes plotted against CG content

This depletion cannot be explained by CG content nor calculated probe specificity (see HybriSeq split probe design in methods) (Fig. 2g-h). We confirmed that this underrepresentation of probes for parts of EIF2S2 and GHITM transcripts is not due to poor probe accessibility or detection efficiency issues by removing RNA from its *in situ* context. We performed *in vitro* bulk hybridization and ligation by extracting total RNA from HEK293 cells, treating RNA with proteinase K to remove protein, hybridizing tiling probes, and capturing RNA with Poly(dT) magnetic beads to enrich mRNA. To rule out cis/trans RNA secondary structures, we included conditions in which RNA devoid of protein was denatured in the presence of probes. We observed close to no representation of probes from the regions that were underrepresented in our single-cell experiments and little difference between native and denatured RNA (Supplementary Fig. 6a). We suspected that these segments of the RNA, which have relatively few probes associated with them, could be missing almost entirely. To determine if this was indeed the case, we examined another dataset that captured full-length polyadenylated RNA sequences. To this end, we analyzed 15 long-read sequencing datasets (Supplementary Table 3) from the Long-read RNA-Seq Genome Annotation Assessment Project (18). We observe little representation of probe-depleted regions in all datasets (Supplementary Figs. 6b-c) and a high fraction of reads truncated before their annotated polyadenylation site (Supplementary Figs. 6b- c), suggesting that for these transcripts, there may be a missing annotation in the reference transcriptome. These results demonstrate that HybriSeq has the potential to reveal intra-transcript biological variabilities.

### Linear Signal Amplification using Multiple Probes

Having demonstrated that the probe-to-probe variability across the same transcript is low with HybriSeq, we reasoned that multiple probes for the same transcript act to linearly amplify the signal to enhance the signal-to-noise ratio (SNR). Thus, we created a simple mathematical model (Supplementary Fig. 7a) by approximating the detection of a specific transcript or probe as a binomial trial (exactly two possible outcomes: detected and not detected). In the case of a scRNAseq measurement, there is only one chance to capture a transcript (Supplementary Fig. 7b), leading to potential “dropout” and excess noise caused by a single priming event. In this scenario, our model predicts that signal scales with the number of transcripts present and the efficiency of capture, the noise is proportional to the square root of the signal, and the SNR is also proportional to the square root of the signal (Supplementary Fig. 7a). Applying this model with the best detection efficiency reported of 45% and an SNR threshold of 2, the lowest number of RNAs reliably detected is 8. With the more typical detection efficiency of 10%, this number is closer to 40 RNAs/cell (Supplementary Fig. 7d). Now for the same model with a linear amplification factor as in HybriSeq (Supplementary Fig. 7c) and an average detection efficiency for a single probe of 20%, a similar or better lower limit of detection can theoretically be accomplished with > 2 probes (Supplementary Fig. 7d).

To confirm our model’s predictions, we subsampled varying numbers of probes for each transcript from our tiling experiment and calculated the signal (UMIs/cell), noise (signal standard deviation), and SNR as a function of the number of probes sampled. Our model predicts that for a given transcript number and capture efficiency, the number of UMIs/transcript will increase in a linear fashion with respect to the number of probes subsampled, the noise will fall off as 1/square root of the number of probes subsampled, and the SNR will increase as a function of the square root of the number of probes subsampled (Fig. 3a). We observed for all probed transcripts that our simple model explains the trends in the signal, noise, and SNR (Figs. 3b-d, Supplementary Fig. 8a-c). For all but one of the transcripts tested (NEFH) (Figs. 3b), we were able to achieve an SNR > 2 with fewer than 6 probes (Figs. 3c-d, Supplementary Fig. 8a-c). In fact, near single-molecule/cell sensitivity can be achieved when > 20 probes are used (e.g. SCAF8 and ARL5B) when summing up all probes (Fig. 3c-d).

**Fig. 3.**
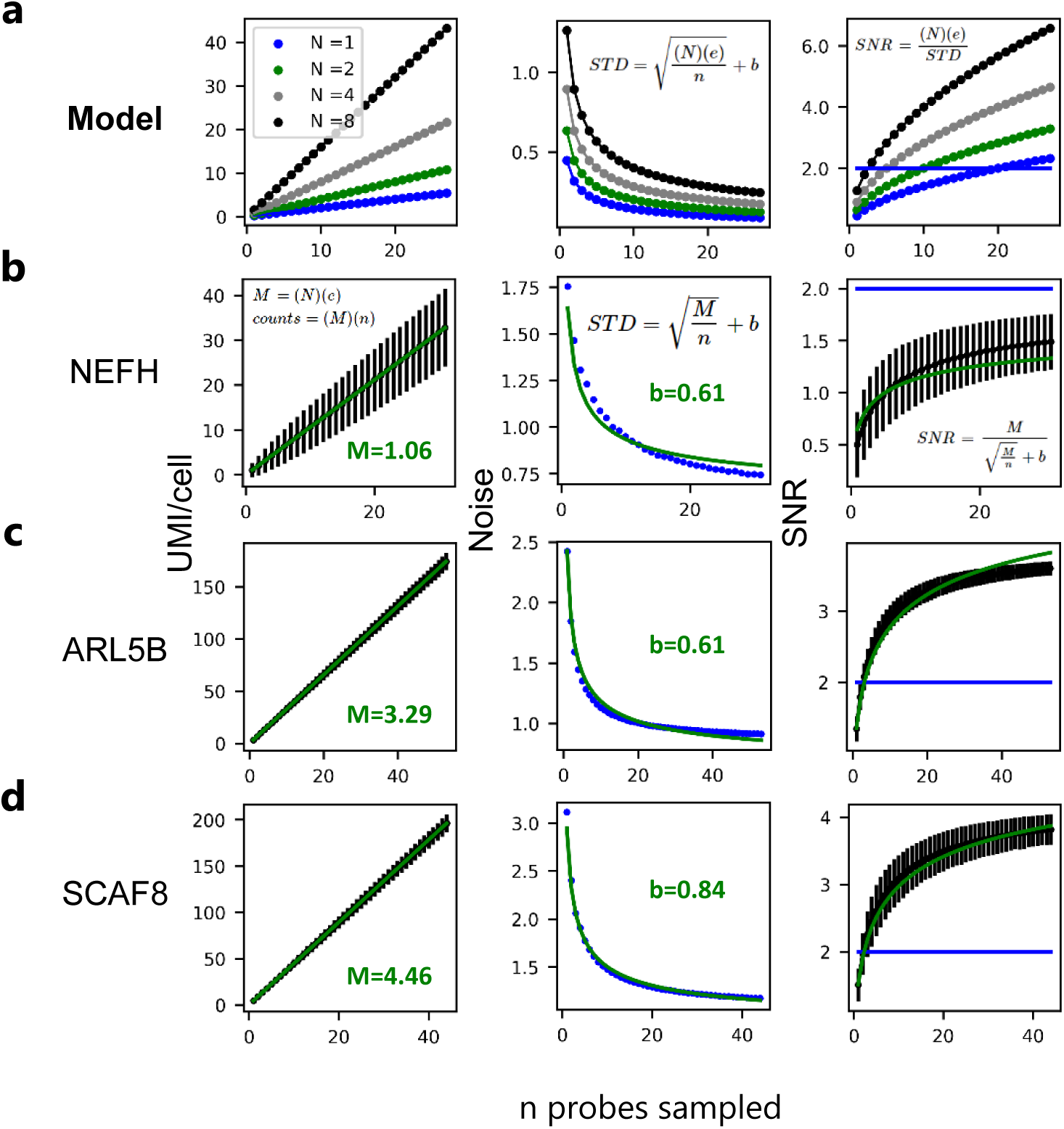
Linear Amplification with HybriSeq. **a.** Left: model of UMIs/cell with n probes, detection efficiency *e* = 0.2, and N transcripts/cell from 1 to 8 transcripts/cell. Middle: model of the measurement standard deviation for UMIs/cell, right: model of the measurement SNR. The blue horizontal line denotes SNR = 2. **b**. Left: Experimentally obtained values for average NEFH UMIs/cell for the sampled number of probes n. Error bars represent the standard deviation associated with each measurement obtained by bootstrap via sampling the unique probes used to calculate average UMIs/cell. The green line represents the fitted linear curve with slope *M*. Middle: Experimentally obtained values for the noise in expression measured with multiple probes *n*. The green line represents the fitted model from (Supplementary Fig. 7A) with the addition of a baseline variance to account for non-measurement-associated heterogeneity. Right: Experimentally obtained values for the measurement SNR measured with multiple probes n. Error bars represent the standard deviation associated with each measurement obtained by bootstrap via sampling the unique probes used to calculate average SNR. The green line represents the fitted model from the middle, and the blue horizontal line denotes SNR = 2. **c.** Same as **b** for ARL5B. **d.** Same as **b** for SCAF8.

### Probing cell-to-cell heterogeneities with HybriSeq

PBMCs contain many cell types important in immune response and have differential transcriptional responses to CD3/CD28-stimulated activation that change over time (19, 20). To demonstrate the ability of HybriSeq to characterize such cell types and response, we constructed a probe set targeting 196 genes known to be important in classifying major cell types in PBMCs, as well as genes known to be differentially expressed in response to CD3/CD28-stimulated activation (Supplementary Table 4). (19, 20). We stimulated PBMCs for 6 and 24 hours and characterized the transcriptional response with HybriSeq. To assess the specificity of this probe set, we again counted the nonspecific ligation events that occurred. We observed that specific ligation events were >22-fold higher than nonspecific ligation events.

After filtering for the number of UMIs and the number of genes captured, we obtained 11,148 single PBMC-targeted RNA profiles. Principal component analysis (PCA) was used for dimensionality reduction on the cell-gene matrix. Cells were clustered in PCA space using the Leiden algorithm, and cell types were assigned to clusters (Fig. 4a) based on marker gene expression (Fig. 4b). 6 hour and 24 hour stimulated cells form distinct separate clusters for all cell types except natural killer (NK) cells when visualized in the UMAP projection (Fig. 4c). For CD4 T cells and B cells we determined genes that were differentially expressed at both 6 and 24 hours as well as genes only differentially expressed at 6 hours or 24 hours (Supplementary Fig. 9). In B cells we observed differential expression of STAT1, IRF1, CD40 and TNF, factors associated with immune activation (21, 22, 23) and inflammatory signaling (24), at both 6 and 24 hours (Fig. 4d). Similarly, in CD4 T cells we observed differential expression of CD69, CD74, IL2RA, and TNF, factors associated with T cell activation (25, 26, 27) and cytokine signaling (24), at both 6 and 24 hours (Fig. 4d).

**Fig. 4.**
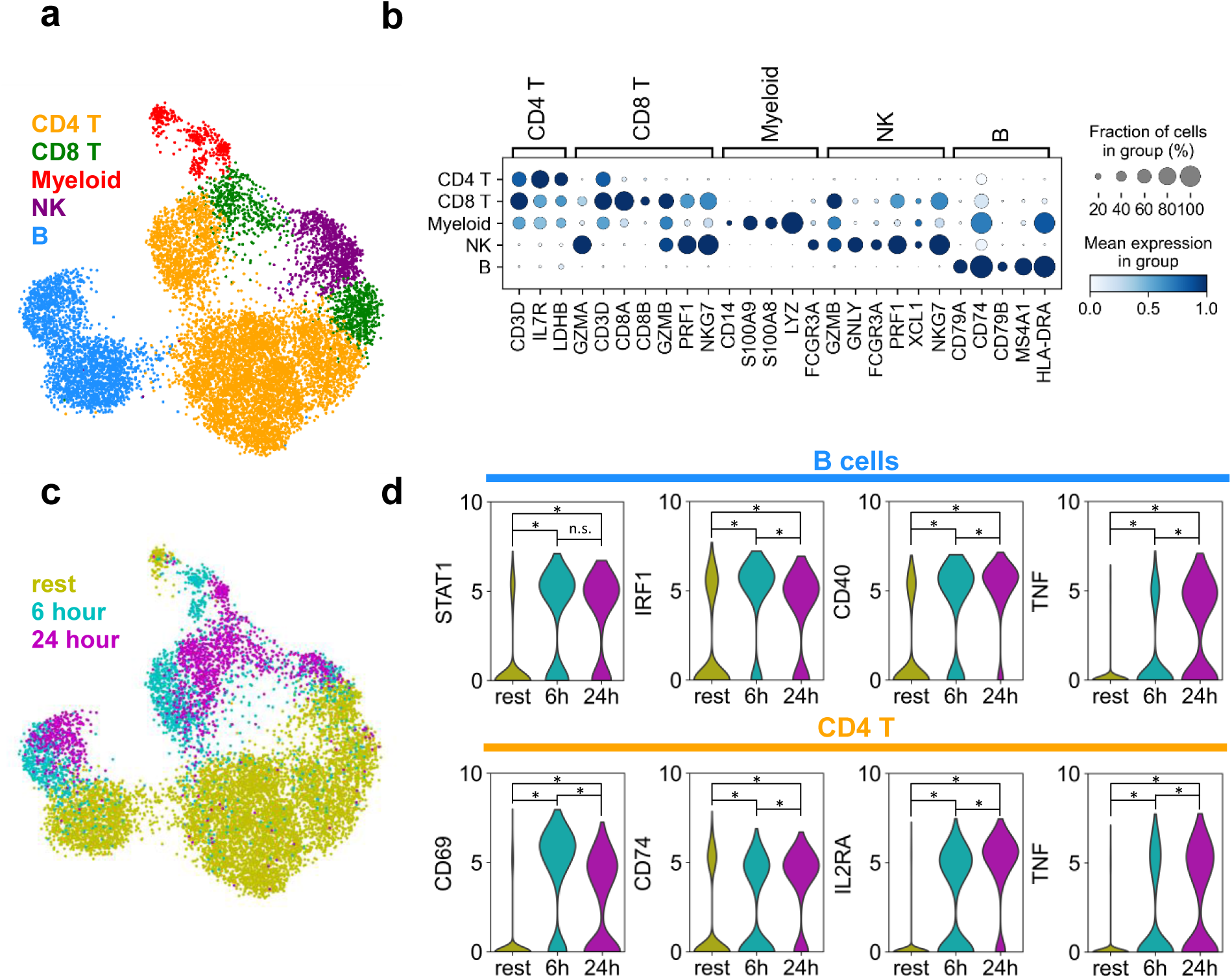
Analyzing PBMC Activation with HybriSeq. **a.** UMAP of 11,148 stimulated and unstimulated PBMCs profiled with probes targeting 196 genes known to be important in classifying major cell types in PBMCs as well as genes known to be differentially expressed in response to CD3/CD28-stimulated activation, colored by cell type. **b.** Dot plot of genes used to determine cell type in **a**. **c.** UMAP same as **a**, colored by the amount of time stimulated with CD3/CD28. Rest cells were not stimulated. **d.** Violin plots of log-normalized expression for select genes that are differentially expressed at both 6 and 24 hours of CD3/CD28 stimulation compared to rest cells for B cells and CD4 T cells. Asterisks denote statistically significant (adjusted p-value <0.01) differential expression between groups. n.s. denotes non-significant.

## Discussion

Here, we present HybriSeq, a probe-based, microfluidics-free method to sensitively profile a set of targeted RNA in single cells. HybriSeq provides a unique set of advantages that overcome current limitations in scRNAseq approaches. First, by utilizing many probes per transcript, HybriSeq offers the ability to confidently detect low-expression transcripts by decreasing the measurement noise. Second, because of the targeted and scalable nature of probe-based split-pool methods, HybriSeq can cost-effectively profile specific biology in many cells by only including probes for transcripts of interest, which greatly increases the efficiency of sequencing and reduces the cost. Finally, HybriSeq utilizes a split-pool approach to label cells with unique cell barcodes, which eliminates the need for microfluidic devices used in other probe-based single-cell RNA profiling methods (10,11,12). This feature allows for the use of cost-effective, off-the-shelf reagents and a simple protocol that is accessible to most users. The cost per cell is approximately 1.4 cents (Supplementary Table 5), excluding sequencing cost, which is similar to the SPLiT-Seq cost of 1-2 cents per cell (13) and much lower than the 10X Genomics Flex approach, which is approximately 10 cents per cell. The unique features of HybriSeq unlock possibilities that were once unattainable with conventional scRNAseq methods. The distinctive features of HybriSeq lie in its ability to accurately quantify RNA expression and intra-transcript variation across diverse transcripts, facilitating the study of cellular transcriptional heterogeneity with heightened sensitivity and resolution.

While powerful in its ability to sensitively detect RNA, the sensitivity of HybriSeq and other probe-based single-cell RNA profiling methods is limited by the length of the RNA molecule being measured, which restricts the total number of probe binding sites. This is the case for all *in situ* hybridization-based approaches and methods utilizing random priming or cDNA fragmentation. For short RNA targets, the number of probes able to hybridize to a transcript could be small even with reduced probe length. A potential workaround to this problem is to use probes with partially overlapped hybridization target regions, as has been utilized in multiplexed FISH methods (9). Moreover, although probe-based methods are efficient in measuring transcript abundance, they are not designed to sequence the RNA molecule itself, thus rendering them inappropriate for detecting RNA sequence variants or modifications. Last, a limitation of HybriSeq is that probe hybridization and cell barcoding require multiple rounds of washes as well as multiple ligation steps. Each of these steps is associated with inefficiency that contributes multiplicatively to decreased sensitivity. Increasing the probe number per transcript could, in some cases, compensate for these inefficiencies.

Our transcript tiling results have shown low probe-to-probe variability for the vast majority of probes. For some transcripts, targeted probes showed lower abundance than those targeting the rest of the transcript, which we conclude is due to the absence of this region of the transcript and not the lack of probe accessibility. Indeed, we confirmed that this absence of probe binding was due to missing annotations. While it is known that the UTR of transcripts can be highly structured and interact with regulatory proteins (28, 29, 30), we were not able to see this impacting probe binding in the transcripts we tested. Considering that certain probes show higher cell-to-cell variabilities compared to other probes targeting the same transcript, this pattern of enrichment/depletion may indeed be indicative of underlying biology pertinent to gene expression regulation and cell-to-cell heterogeneity. In the case of transcripts with alternative splicing, such analysis can be performed by including probes for introns and across splicing junctions, showcasing the advantage of non-3’-biased detection in HybriSeq. Furthermore, investigation into this phenomenon will also yield useful insights into probe design for FISH-based spatial transcriptomic approaches, which rely on hybridization to make measurements.

## Methods

### HybriSeq split probe design

HybriSeq ssDNA probes are composed of five regions split into two probes as follows from 5’ to 3’:

1. (Left probe) 20 nt priming region, which is a partial Illumina Nextara read 2 or a different universal priming region (tiling probes).
2. (Left probe) 30 nt left probe targeting region.
3. (Right probe) 30 nt right probe targeting region. The first two bases are either A or T. SplintR ligase has higher efficiency when C or G are not in the first two bases of the ligation site.
4. (Right probe) 8 nt random UMI sequence
5. (Right probe) 15-20 nt round 1 ligation handle

A probe design pipeline was adapted from Moffitt et al. (31), with minor changes. For calculating gene and isoform level specificity of probes, our pipeline only considers the center 30 nt of the targeting region (last 15 nt of the left probe + first 15 nt of the right probe) and does not directly consider melting temperature as a parameter when selecting probes, but considers CG content. To calculate the probe specificity index, our approach differs from Moffitt et al. (31) in that we do not consider transcript abundance and set all transcripts’ abundance equal to 1. Additionally, we use a 15-nt window as opposed to a 17-nt window when calculating the penalty lookup tables.

Probe specificity (Fig. 2g) measures how often a given 15nt sequence within a probe appears in the entire transcriptome. Theoretical probe specificity is calculated as the proportion of that probe’s occurrences within all isoforms of the target gene compared to its occurrences across all transcripts in the transcriptome.

Probes were obtained from IDT (Integrated DNA Technologies) in the 50 nmole oPools format or individually as single probes ordered as DNA oligos (Supplementary Tables 1, 2, 4, 10).

Probe design and all analysis use human reference genome GRCh38 and mouse reference genome GRCm39.

Right side probes were 5’ phosphorated with T4 Polynucleotide Kinase (New England Biolabs - NEB ref M0201S). Probes were then column cleaned with ssDNA/RNA Clean & Concentrator (Zymo D7010) and quantified. Left-side probes were added at an equal molar concentration and used in hybridization.

### Cell culture

HEK293 cells were originally obtained from the UCSF Cell Culture Facility. HeLa (CCL-2) cells were originally obtained from ATCC. Cell line stocks were cultured in DMEM + 10% FBS and 1% Penicillin-Streptomycin. Cells were washed twice with 1X PBS, then detached by incubating 2-5 minutes at room temperature with 3 mL of 0.25% Trypsin. Once cells were detached, they were added to 7 mL of media with 10% FBS. In cell mixing experiments, cells were combined at the desired concentrations at this step. Cell lines were not tested for mycoplasma contamination and were not authenticated.

Do-11-10 cells (Sigma-Aldrich 85082301) were cultured in DMEM + 10% FBS and 1% Penicillin-Streptomycin. Suspended cells were collected in a 15 mL conical tube and pelleted for 3 minutes at 300g. Cells were then resuspended in 10 mL of PBS. In cell mixing experiments, cells were combined at the desired concentrations at this step.

For PBMC experiments, cryopreserved human peripheral blood mononuclear cells from a healthy donor were purchased from Cytologics. PBMCs were thawed in a 37 °C water bath and transferred to a conical vial containing 10 mL of pre-warmed RPMI-1640 (Gibco) media + 1% Penicillin-Streptomycin. Cells were pelleted for 10 minutes at 300g and resuspended in pre-warmed RPMI-1640 + 1% Penicillin-Streptomycin + 10% FBS and plated into an ultra-low attachment 6-well plate (Costar) at a final density of 1 x 10^6^ cells/mL. Cells were incubated overnight at 37 °C, 5% CO2. Cell supernatant was removed and placed in a 15 mL conical tube and 1 mL TrypLE was added to the plate and incubated a 37°C, 5% CO2 for 4 minutes to lift adherent cells. TrypLE was removed and added to the 15 mL conical tube containing the cell supernatant. Cells were pelleted 300g for 5 minutes. Cells were resuspended in pre-warmed RPMI-1640 + 1% Penicillin-Streptomycin + 10% FBS and plated in a 96 well round bottom ultra-low attachment plate (Corning) at a final density of 2 x 10^6^ cells/mL. ImmunoCult Human CD3/CD28 T Cell Activator (STEMCELL Technologies) was added according to manufacturer’s directions for 6h, 24h, or not added (rest). To retrieve PBMCs to profile with HybriSeq, cell supernatant was removed and placed in a 1.5 mL conical tube and 0.1 mL TrypLE was added to the plate and incubated a 37°C, 5% CO2 for 4 minutes to lift adherent cells. TrypLE was removed and added to the 1.5 mL conical tube containing the cell supernatant.

### Fixation

Cells were centrifuged for 3 minutes at 500 g at 4 °C. Cells were washed in 1 mL of 1X PBS. The cells were then passed through a 40 μm strainer into a 15 mL falcon tube and counted. Cells were centrifuged for 3 minutes at 500 g at 4°C. Cells were resuspended in 0.5 million cells/mL of 4% freshly prepared paraformaldehyde solution in 1X PBS. Cells were fixed for 30 minutes at room temperature under gentle agitation. Cells were centrifuged for 3 minutes at 500 g at 4 °C and washed 2 times in 1X PBS. The cells were then passed through a 40 μm strainer into a 15 mL falcon tube and counted.

### Hybridization & Ligation

Cells were resuspended in hybridization buffer (30% formamide, 1% BSA, 0.5% Tween 20, 2X SSC, 40U/mL RNasin) for 10 minutes at 37 °C under gentle agitation. Cells were centrifuged for 3 minutes at 500g at 4°C. Cells were resuspended in a hybridization buffer with probes at 10nM/probe. Cells were incubated at 37 °C for 18-24 hours with gentle agitation. Cells were then washed in wash buffer (20% formamide, 0.5% Tween 20, 4X SSC, 40 U/mL RNasin) two times at 37 °C for 5 minutes. Cells were washed in ligation buffer (1X T4 DNA Ligase Reaction Buffer (NEB ref B0202S), 0.4 mM ATP, 40 U/mL RNasin) and then resuspended in ligation buffer plus 2 µM SplintR Ligase (NEB ref M0375S). Cells were incubated for 1 hour at 37 °C with gentle agitation.

### RNA Isolation

Suspended cells were washed in 1 mL of 1X PBS. ∼5 million cells were centrifuged for 3 minutes at 500 g at 4 °C and total RNA was extracted with RNeasy Mini kit (Qiagen ref 74104) according to the manufacturer’s protocol. Total RNA was then treated with DNase I (NEB ref M0303L) by combining 10µg of total RNA in 1X DNase I reaction buffer (NEB ref M0303L) with 2 units of DNase I (NEB ref M0303S) and heating at 37 °C for 15 minutes. For *in vitro* hybridization experiments 8U of Proteinase K (NEB ref P8107S) was also included in the reaction. DNA-depleted RNA was extracted with an RNeasy Mini kit (Qiagen ref 74104) according to the manufacturer’s protocol.

### *In vitro* Hybridization, Ligation, and mRNA Capture

1µg of RNA was diluted in *in vitro* hybridization buffer (final concentration: 30% formamide, 2X SSC, 40U/ml RNasin) with probes (10nM/probe) to a total volume of 25µL. For the denatured condition, RNA was heated at 90 °C for 5 minutes before being incubated at 37 °C for 18-24 hours. For the native condition, RNA was incubated at 37 °C for 18-24 hours. After hybridization, 0.2mg of Oligo d(T)25 Magnetic Beads (NEB ref S1419S) in 75µL of 20X SSC were added to the RNA. RNA was incubated at 37 °C for 5 minutes before being cooled to 4 °C at 0.1 °C/second to anneal RNA to beads. Tubes containing the bead/RNA hybrids were placed against a strong magnet for 2 minutes until the solution was clear. Beads were washed two times in 20X SSC to remove unbound RNA, and wash solution was removed. 50µL of 1X DNA ligation buffer plus 2 µM SplintR Ligase was added to the Bead/RNA hybrids and incubated for 1 hour at 37 °C to ligate adjacent RNA-bound probes. 150µL of 20X SSC was added to the ligation mix and incubated for 5 minutes at 25 °C. Tubes containing the bead/RNA hybrids were placed against a strong magnet for 2 minutes until the solution was clear. Bead/RNA hybrids were resuspended in 1X RNase H buffer containing 5 units of RNase H (NEB ref M0297S) and incubated at for 1 hour at 37 °C to release probes into solution. Tubes containing the mixture were placed against a strong magnet for 2 minutes until the solution was clear, and the solution was collected for sequencing library preparation.

### Preparing Oligos for Ligations

The first and second barcoding steps consist of a ligation reaction. Each round uses a different set of 96 well barcoding plates. Ligation rounds have a universal linker (Supplementary Table 6) strand with partial complementarity to a second strand containing the unique well-specific barcode sequence added to each well (Supplementary Table 7, 8). These strands were annealed together prior to barcoding to create a DNA molecule with three domains: a 15 nt 5’ overhang that is complementary to the 15 nt 3’ overhang present on the right-side probe, a well-specific barcode sequence, and a 15 nt 3’ overhang complementary to the 5’ overhang present on the next barcode molecule to be subsequently ligated. For the experiments to produce Fig. 1e, an additional five random nucleotides were included on the 3’ side of the well-specific barcode sequence. For the second-round barcodes, the 3’ overhang acts as a universal priming region to which the third round well-specific primer can anneal and extend in a PCR. Barcode strands for the ligation rounds are added to 96 well plates and their 5’ ends phosphorylated with T4 Polynucleotide Kinase (NEB ref M0201S). After 5’ phosphorylation, equal molar amounts of linker strand are added to each well making the final concertation 5.4 µM. Oligos for ligation are annealed by heating plates to 95 °C for 2 minutes and cooling down to 20 °C at a rate of –0.1 °C per second. For ligation reactions, 2.31 µL of barcode/linker oligos are added to 96 well plates to which cells can be added.

### Cell barcoding

After probe ligation cells were counted and added to the ligase buffer (1X T4 DNA Ligase Reaction Buffer (NEB ref B0202S), 0.4 mM ATP, 40 U/mL RNasin, 0.5% Tween 20, 1% BSA, 200,000 U/mL T4 ligase) so that the final cell concentration was 22,000 cells/mL. Cells were passed through a 40 μm strainer. 22.69 µL of cells (500 cells) in ligase buffer were added to each well of a 96-well protein low bind plate, which had 2.31 µL of barcode 1 and linker 1 oligos already in each well. For the PBMC experiment, each stimulation condition was assigned to a specific set of wells, and this information was recorded. The first barcode was then used to identify the corresponding stimulation condition. Columns 1-4: Rest, columns 5-8: 6 hours, and columns 9-12: 24 hours. Cells were mixed by gently pipetting up and down. Plates were sealed and incubated at 25 °C for 2 hours. 2 µL of 62.5 µM quenching oligo 1 (Supplementary table 6) were added to each well and mixed by pipetting. Plates were sealed and incubated at 25 °C for 30 minutes. 25 µL of barcode wash buffer (50 mM EDTA, 0.5% Tween 20) was added to each well and incubated for 10 minutes. Cells from all 96 wells were pooled into a single 5ml low-bind Eppendorf tube. Cells were centrifuged for 3 minutes at 500 g at 4 °C. Cells were washed two times in barcode wash buffer (+5 µM quenching oligo 1) and then washed in ligase buffer (+5 µM quenching oligo 1, -T4 ligase). Cells were resuspended in ligase buffer (+5 µM quenching oligo 1) so that the final cell concentration was 22,000 cells/mL and passed through a 40 μm strainer. 22.69 µL of cells in ligase buffer (+5 µM quenching oligo 1) were added to each well of a 96-well protein low bind plate, which had 2.31 µL of barcode 2 and linker 2 oligos already in each well. Cells were mixed by gently pipetting up and down. Plates were sealed and incubated at 25 °C for 2 hours. 2 µL of 62.5 µM quenching oligo 2 were added to each well and mixed by pipetting. Plates were sealed and incubated at 25 °C for 30 minutes. 25 µL of barcode wash buffer was added to each well and incubated for 10 minutes. Cells from all 96 wells were pooled into a single 5ml low-bind Eppendorf tube. Cells were centrifuged for 3 minutes at 500 g at 4 °C. Cells were washed two times in barcode wash buffer (+5 µM each of quenching oligo 1 & 2) and then resuspended in ice-cold 1X ThermoPol reaction buffer (NEB ref B9004S). Cells were passed through a 40μm strainer and counted. Cell concentration was normalized to 23,000 cells/mL in a cold ThermoPol reaction buffer. 460 cells/well were dispensed into many 200µl PCR tubes or 96 well PCR plates. 20µL of PCR solution (1X KAPA HiFi HotStart ReadyMix (final concentration) and forward primer) with well-specific round 3 reverse primers (Supplementary Table 9) were added to each well, so that the final concentration of each primer was 0.4 µM. PCR thermocycling was performed as follows: 95 °C for 30 seconds, then 20 cycles at 95 °C for 30 seconds, 55 °C for 30 seconds, 72 °C for 30 seconds, followed by a final extension at 72 °C for 30 seconds. For the probe tiling experiment 48 wells were used in each barcoding round with the final round 3 PCR barcoding reaction containing 115 cells/well.

### Library Preparation for Sequencing, Single Cell

Round 3 PCR reactions were centrifuged at 500g for 1 minute to pellet cells. All round 3 PCR reaction solution was removed, pooled, and column purified with the Zymo DNA clean & concentrator kit (Zymo 11-305). Purified libraries were analyzed on an Agilent TapeStation Systems (D1000 kit) to check for the correct DNA size. If the predominate band was the correct size (252 ± 2 bp or 232 ± 2 bp depending if the left probe included a partial read 2 priming sequence) and was < 90% of the library the purified PCR product was run on a 3% agarose Tris-acetate-EDTA electrophoresis gel (200V 20 minutes) and the correct size band was cut out and extracted from the agarose with the Zymo Gel recovery kit (Zymo D4002). We observe that libraries that contained left probes containing the non-read 2 priming regions produced some nonspecific amplification, requiring size selection purification. The purified pooled round 3 DNA product was placed into a final limited-cycle PCR to add Illumina sequencing adaptors. The adapter addition PCR reaction was as follows: 0.5 ng DNA from pooled round 3 PCR product, 0.4 µM P7 forward primer, 0.4 µM P5 reverse primer, and 1X KAPA HiFi HotStart ReadyMix. PCR thermocycling was performed as follows: 95 °C for 30 seconds, then 10 cycles at 95 °C for 30 seconds, 55 °C for 30 seconds, 72 °C for 30 seconds, followed by a final extension at 72 °C for 30 seconds. The PCR reaction was removed and purified with a 0.8X ratio of AMPure XP beads to generate an Illumina-compatible sequencing library.

### Library Preparation for Sequencing, Bulk HybriSeq

For bulk *in vitro* HybriSeq experiments, 1ul of RNase H digestion product was loaded into a 25 µL 1X KAPA HiFi HotStart ReadyMix PCR reaction with 0.3 µM primers containing partial Illumina sequencing adapters. PCR thermocycling was performed as follows: 95 °C for 30 seconds, then 15 cycles at 95 °C for 30 seconds, 55 °C for 30 seconds, 72 °C for 30 seconds, followed by a final extension at 72 °C for 30 seconds. 1 µL of this unpurified PCR reaction was then added to a similar PCR reaction containing the primers for the addition of the full Illumina sequencing adaptors. PCR thermocycling was performed as follows: 95 °C for 30 seconds, then 8 cycles at 95 °C for 30 seconds, 55 °C for 30 seconds, 72 °C for 30 seconds, followed by a final extension at 72 °C for 30 seconds. The PCR reaction was removed and purified with a 1.3X ratio of AMPure XP beads to generate an Illumina-compatible sequencing library.

### RNA-seq Library Preparation, Sequencing, and Analysis

Library preparation and sequencing were performed by the QB3-Berkeley Genomics core labs. Total RNA quality as well as poly-dT enriched mRNA quality were assessed on an Agilent 2100 Bioanalyzer. Libraries were prepared using the KAPA mRNA Hyper Prep kit (Roche KK8581). Truncated universal stub adapters were ligated to cDNA fragments, which were then extended via 10 cycles of PCR using unique dual-indexing primers into full-length Illumina adapters.

Library quality was checked on an AATI (now Agilent) Fragment Analyzer. Library molarity was measured via quantitative PCR with the KAPA Library Quantification Kit (Roche KK4824) on a BioRad CFX Connect thermal cycler. Libraries were then pooled by molarity and sequenced on an Illumina NovaSeq X, 25B flowcell for 2 x 150 cycles, targeting at least 25M reads per sample. Fastq files were generated and demultiplexed using Illumina BCL Convert and default settings, on a server running CentOS Linux.

Reads from the generated Fastq files were mapped to the mouse transcriptome (GRCm39) and transcripts per million (TPM) quantified using Salmon (32). The abundance of binding sites for each probe in the HybriSeq experiments was calculated by summing the expression of all transcripts to which the probe could fully hybridize, accounting for multiple binding sites on the same transcript. Probe counts and transcript TPM values for the same gene were then averaged and plotted against each other.

### Illumina Sequencing

15 pM libraries were sequenced on a MiSeq (Illumina) using a 150-nucleotide (nt) V3 kit in paired-end format. Read 1 (75 nt) covered the cell barcode, and read 2 (75 nt) covered the probe and UMI.

### RNase H specificity

After non-split probe hybridization and washing, cells were resuspended in RNase H reaction buffer containing 20 U/mL of RNase H enzyme (NEB M0297S). Cells were incubated for one hour at 37 °C with gentle agitation. Released probes were quantified with sequencing or qPCR.

### qPCR

qPCR was performed on probes or barcoded probes (Supplementary table 10) released from cells via RNase H release or heat release. Cells and released probes were centrifuged to pellet cells, the supernatant was purified with spin columns, and DNA was eluted in 20ul of water (Zymo ssDNA/RNA clean & concentrator). 1 µL of purified sample was loaded into each 20 µL reaction of a qPCR with 0.3 µM primers (Supplementary table 10) according to the manufacturer’s instructions using Maxima SYBR Green qPCR Master Mix (Thermo Fisher Ref K0222). Thermocycling and measurements were performed on a QuantStudio™ 5 System with the following temperatures: 95 °C for 60 seconds, 45 cycles of 95 °C for 15 seconds and 60 °C for 30 seconds (1.6 °C/second ramp rate) in which the fluorescence was recorded at the end of each cycle. QuantStudio™ Design and Analysis Software v1.4.1 was used to analyze fluorescence signal and calculate CT values. A standard curve was made by running a dilution series of the target oligo (ordered from IDT), and the CT values from this were used as the standard curve from which the concentrations of target oligos in the sample were determined. To measure the % ligated barcode in supplementary Fig. 3, the concentration of ligated species (right side probe + barcode 1) was compared to the concentration of just the right-side probe present in the reaction.

### HybriSeq Computational Pipeline

We constructed a pipeline to analyze HybriSeq data by taking raw sequencing reads and constructing a count matrix (counts per probe per cell). Briefly, we identify real barcodes, identify probe-targeting regions with correct ligation, remove duplicates using UMIs, and filter out reads not containing barcodes or probe-targeting regions. Detailed key steps were as follows:

1. To determine the location of each 7-base barcode sequence on read 1, we calculated the reverse complement of read 1, searched for the location of regions that should be common to all reads, and extracted the 7-base barcode with these locations. More specifically, to determine the location of barcode 1, we searched the first 25 bases of read 1 for the sequence *ATTCG*, common to all barcode 1 oligos, and used the 9 bases after to determine barcode 1. To determine the location of barcode 2, we searched from bases 35 to 56 of read 1 for the sequence *TGCTTGAG,* common to all barcode 2 oligos, and used the 9 bases after to determine barcode 2. To determine the location of barcode 3, we searched from base 57 to the end of read 1 for the sequence *GTTTCG,* common to all barcode 3 oligos, and used the 7 bases after to determine barcode 3. If the sequences *ATTCG, TGCTTGAG,* or *GTTTCG* were not found in their respective search windows, the read was excluded from analysis.
2. To determine the unique cell barcode, whitelists of each round of unique barcode sequences were constructed. We search the bases extracted from step 1 for 7 base stretches that were in the whitelists. If no match or multiple matches are found, reads were excluded. From this, a unique cell barcode was constructed.
3. To determine the targeting region from read 2 for the probe tiling experiments, a whitelist of probes was constructed, including probe sequences within a Hamming distance of two. Left-side probe targeting regions were determined by taking the first 30 bases of read 2 and comparing them to the probe whitelist. Similarly, right-side probe targeting regions were determined by taking the next 30 bases of read 2 and comparing them to the probe whitelist. Reads containing targeting regions not predicted to be adjacent were excluded. For all other experiments, the targeting region was determined by aligning read 2 to a custom index using bowtie2 (33) with the parameters: -L 30, and reads trimmed so that only the 60 bases of the ligated probe are aligned. The custom index included every combination of left- and right-side probes so that nonspecific ligation events could be calculated (Supplementary Fig. 4a). The 8 bp simple UMI included on the right-side probe was extracted for both approaches.
4. We constructed a data frame of reads that included the unique cell barcode, probe targeting region, and simple 8bp UMI. We then collapsed duplicate reads by considering a combined UMI that contained the 8 bp simple UMI, the unique cell barcode, and the identity of the probe targeting region.
5. We generated a count table of UMIs per probe per unique cell barcode or UMIs per gene per unique cell barcode.
6. To determine which unique cell barcodes were associated with real cells, a threshold for UMIs/cell was calculated by taking 10% of the 99^th^ percentile of the top set of unique cell barcodes or visually setting the threshold at the first knee of the cell rank - UMI plot. We note that when only considering lowly or highly variably expressed transcripts, the inclusion of probes targeting moderately and stably expressed transcripts can help set a threshold.
7. The Scanpy library (34) in Python was used for all standard single-cell analyses.

For PBMC experiments, cells were clustered using the Leiden algorithm on the PCA embedding, and cell types were assigned to clusters based on expression of marker genes. Differential expression of genes in every cell type across stimulation conditions was determined using the Scanpy (34) function rank_genes_groups using the Wilcoxon method and Benjamini-Hochberg correction. For each cell type, a comparison was made between the unstimulated cells (rest) and the 6-hour or 24-hour CD3/CD28 stimulated cells. A gene was considered differentially expressed in a given cell population if the Benjamini-Hochberg corrected p-value was less than 0.01.

### Mixing/ Single-Cell Purity Experiment

A mixture of HEK293 cells and Do-11-10 cells was subjected to the standard HybriSeq protocol. Equal concentrations of each cell line were mixed before the fixation step. Probes (2-3 probes per transcript) targeting 95 human cell cycle-associated transcripts and 124 mouse cell cycle-associated transcripts were added to the cell mixture during the hybridization step.

### Probe Tiling Analysis

Each transcript was analyzed independently, only considering probes targeting that transcript. Probe counts for each cell were normalized so that the total sum of all normalized counts in each cell was equal to the median UMIs/cell of the cell population. This was done to account for differences in expression levels between cells. The average relative counts were taken for each probe and plotted as a trace for all cells. The standard deviation was calculated for each probe. For bulk *in vitro* and *in situ* tiling experiments, each transcript was analyzed independently, only considering probes targeting that transcript. Probe counts were normalized so that the total sum of all normalized counts was equal to 200. The standard deviation was calculated for each probe from three independent experiments.

### Long-read RNA-Seq Genome Annotation Assessment Project (LRGASP) Data Analysis

15 datasets were selected from the LRGASP by querying untreated/unperturbed polyadenylated long-read human cell line RNA sequencing data generated using the Pacific Biosciences Sequel system, which was replicated and had sufficient sequencing depth (Supplementary Table 3).

Binary Alignment Map (BAM) files, excluding unfiltered BAM files, were retrieved from ENCODE (Supplementary Table 3). Sequences that mapped to chr20:34088309-34112243 (EIF2S2) and chr10:84139482-84154841 (GHITM) were extracted from BAM files using Samtools. For each annotated transcript sequence (EIF2S2: ENST00000374980.3, GHITM: ENST00000372134.6), a list of every 20bp subsequence was created. The number of occurrences of each 20bp subsequence in each sample’s extracted alignments was calculated for both transcripts separately and normalized to the total number of alignments in each dataset to generate the fraction of reads which contain each 20bp subsequence. Standard deviation was calculated across the 15 datasets.

### Measurement Variability Model and Simulation

To model measurement noise associated with sampling a specific transcript in a cell we started off by making a few assumptions.

- Sampling of a transcript in a cell can be modeled with a Poisson distribution.
- The probability of capturing a transcript or probe is the same for all probes targeting the same transcript or priming events.
- The background signal from random probe ligation is minimal and can be assumed to be negligible.
- Probe binding to a transcript does not influence different probes binding the same transcript.
- Probes are hybridized at a saturating concentration.
- The underlying cell-cell heterogeneity can be modeled as a constant value of standard deviation and is not dependent on the number of probes used.
- All cells have the same efficiency of detection for the same transcript. To model single cell transcript measurement variability:

Let N be the number of specific transcripts in a cell, n be the number of detection chances per transcript in a cell, e be the efficiency at which n is successfully detected, and C be the number of counts or UMIs for a specific transcript. If we assume that N is Poisson, the variability associated with counts C is equal to the mean of C and we define measurement noise as the standard deviation of the measurement C:

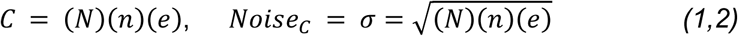

Taking the ratio of the counts C to the noise associated with C we get the signal to noise ratio (SNR)

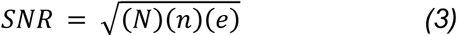

For a population of cells, C will scale linearly with n. If we define expression, M, as C normalized to the number of probes used to make the measurement, expression is given by:

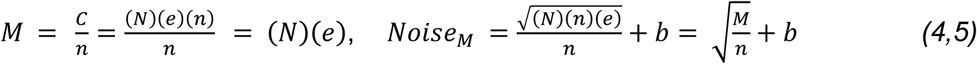

Here we assume that that the contribution to noise in the expression measurement from biology is independent of the number of probes used to make the measurements and can be defined as constant b.

The expression SNR is then given by the ratio of M to Noise associated with M:

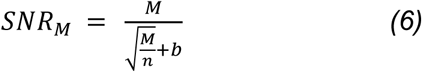

For the simulations in Fig. 3, we experimentally determine M by taking the slope of the line fit to the UMI/cell – probe number plot. This slope is the number of counts you would expect to gain for each additional probe included in the analysis.

We then non-linearly used least squares to fit the function for Noise of M to the standard deviation of M as a function of the number of probes used to make the measurement, keeping M constant from the experimentally determined M and only fitting the model by optimizing b.

### Calculation of signal, standard deviation, and SNR for multiple probes

Total probe counts for each cell were normalized so that the total sum of all normalized counts in each cell was equal to the total median UMIs/cell of the cell population. This was done to account for differences in expression levels between cells, as the goal is to gain an understanding of the measurement-associated variation and not necessarily the underlying inherent biological variation. To calculate the average signal or counts, for each number of probes considered (n), a random set of probes was chosen with replacement, and the average number of UMIs/cell was calculated along with a standard deviation across cells for each n. To calculate the SNR, the ratio of average expression (UMIs/cell/n) to the standard deviation of average expression was calculated for all n. This procedure was repeated 10,000 times, randomly sampling the set of probes used to make the measurement, and the average and standard deviations of these calculations were plotted.

### Statistics and Reproducibility

For Fig.1 e-f, the Pearson correlation coefficient and associated p-values were calculated using the linregress function in the Python library SciPy. For probe tiling experiments, the standard deviation and average were calculated for each probe across all cells profiled. For bulk *in vitro* and *in situ* tiling experiments, each transcript was analyzed independently, only considering probes targeting that transcript. Probe counts were normalized so that the total sum of all normalized counts was equal to 200. The standard deviation was calculated for each probe from three independent experiments. To determine differentially expressed genes in the PBMC experiment, the Scanpy (34) function rank_genes_groups was used with the Wilcoxon method and Benjamini-Hochberg correction. A gene was considered differentially expressed if the Benjamini-Hochberg corrected p-value was less than 0.01.

## Acknowledgements

We thank Eric Chow for the valuable discussions. This work was partially supported by the National Institutes of Health [DK127421 to B.H.]. Funding for open access charge: National Institutes of Health. B.M. is a MacMillan Family Foundation Awardee of the Life Sciences Research Foundation. Q.Z. is supported by a Cancer Research Institute Immuno-Informatics Postdoctoral Fellowship (CRI5054). B.H. and Z.J.G are Chan Zuckerberg Biohub San Francisco Investigators.

## Author Contributions

D.F. performed the experiments, analyzed the data, and contributed to the interpretation of the results. B.H. conceived the project. D.B. and S.Z. conceived and designed the proof-of-concept experiments. B.M. and Q.Z. conceived and performed part of the PBMC experiments. Z.J.G. contributed to project discussions and direction. B.H. and D.F wrote the manuscript.

## Competing Interests

The authors declare no conflict of interest.

## Data Availability

A public git repository is available at https://github.com/BoHuangLab/HybriSeq containing source code and a minimum analytical dataset sufficient to reproduce figures in the main text. This repository is also available through Figshare (https://doi.org/10.6084/m9.figshare.23882193.v2). The fastq files and processed data are available through GEO (GSE292322).

## Notes

### Competing Interest Statement

The authors have declared no competing interest.

### Summary of Updates

Enhanced Specificity Demonstration: Comparative data between split and non-split FLNA probes targeting the same transcript region was added, confirming increased specificity with split probes (Supplementary Fig. 2b-c). Improved Biological Validation: Cell cycle gene experiments were replaced with analysis of peripheral blood mononuclear cells (PBMCs), profiling approximately 11,000 cells to demonstrate the method's utility in a heterogeneous biological system. Scale and Robustness: Additional experiments including a mouse-human cell mixing experiment (~4,000 cells) were conducted to demonstrate scalability and minimal cell mixing (1.8%). Correlation with Bulk RNA-seq: An improved correlation of 0.90 between HybriSeq and bulk RNA-seq in DO-11-10 cells was reported, consistent with other probe-based single-cell approaches.

## References

1. Osumi-Sutherland, D., Xu, C., Keays, M., Levine, A.P., Kharchenko, P.V., Regev, A., Lein, E., Teichmann. S.A. Cell type ontologies of the human cell atlas. Nat Cell Biol. 11, 1129–1135 (2021). 10.1038/s41556-021-00787-7

2. Macosko, E.Z. et al. Highly Parallel Genome-wide Expression Profiling of Individual Cells Using Nanoliter Droplets. Cell. 161(5), 1202–1214 (2015). 10.1016/j.cell.2015.05.002

3. Klein, A.F. et al. Droplet Barcoding for Single-Cell Transcriptomics Applied to Embryonic Stem Cells. Cell. 161(5), 1187–1201 (2015). 10.1016/j.cell.2015.04.044

4. Wulf, M.G., Maguire, S., Humbert, P., Dai, N., Bei, Y., Nichols, N.M., Corrêa, I.R., Guan, S. Non-templated addition and template switching by Moloney murine leukemia virus (MMLV)- based reverse transcriptases co-occur and compete with each other. J BIOL CHEM. 48, 18220–18231 (2019). 10.1074/jbc.RA119.010676

5. Zhang, X., Li, T., Liu, F., Chen, Y., Yao, J., Li, Z., Huang, Y., Wang, J. Comparative analysis of droplet-based ultra-high-throughput single-cell RNA-seq. Systems. Mol. Cell. 73,1,130–142 (2019). 10.1016/j.molcel.2018.10.020

6. Bagnoli, J.W., Ziegenhain, C., Janjic, A., Wange, L.E., Vieth, B., Parekh, S., Geuder, J., Hellmann, I. & Enard, W. Sensitive and powerful single-cell RNA sequencing using mcSCRB-seq. Nat. Commun. 9, 2937 (2018). 10.1038/s41467-018-05347-6

7. Svensson, V., Natarajan, K.N., Ly, L., Miragaia, R.J., Labalette,C., Macaulay, I.C., Cvejic, A., & Teichmann, S.A. Power analysis of single-cell RNA-sequencing experiments. Nat. Methods. 14, 381–387(2017). 10.1038/nmeth.4220

8. Mereu, E., Lafzi, A., Moutinho, C., Ziegenhain, C., McCarthy, D.J., Álvarez-Varela, A., Batlle, E., Sagar, N., Gruen, D., Lau, J.K. and Boutet, S.C. Benchmarking single-cell RNA-sequencing protocols for cell atlas projects. Nature biotechnology 38(6), pp.747–755 (2020). 10.1038/s41587-020-0469-4

9. Wang, G., Moffitt, J.R. and Zhuang, X. Multiplexed imaging of high-density libraries of RNAs with MERFISH and expansion microscopy. Scientific reports 8(1), p.4847 (2018). 10.1038/s41598-018-22297-7

10. Marshall, J.L., Doughty, B.R., Subramanian, V., Guckelberger, P., Wang, Q., Chen, L.M., Rodriques, S.G., Zhang, K., Fulco, C.P., Nasser, J. and Grinkevich, E.J. HyPR-seq: single-cell quantification of chosen RNAs via hybridization and sequencing of DNA probes. Proceedings of the National Academy of Sciences 117(52), pp.33404–33413 (2020). 10.1073/pnas.201073811

11. McNulty, R., Sritharan, D., Pahng, S.H., Meisch, J.P., Liu, S., Brennan, M.A., Saxer, G., Hormoz, S. and Rosenthal, A.Z. Probe-based bacterial single-cell RNA sequencing predicts toxin regulation. Nature Microbiology 8(5), pp.934–945 (2023). 10.1038/s41564-023-01348-4

12. Janesick, A., Shelansky, R., Gottscho, A.D. et al. High resolution mapping of the tumor microenvironment using integrated single-cell, spatial and in situ analysis. High resolution mapping of the breast cancer tumor microenvironment using integrated single cell, spatial and in situ analysis of FFPE tissue. Nat Commun 14, 8353 (2023). 10.1038/s41467-023-43458-x

13. Rosenberg, A.B., Roco, C.M., Muscat, R.A., Kuchina, A., Sample, P., Yao, Z., Graybuck, L.T., Peeler, D.J., Mukherjee, S., Chen, W. and Pun, S.H. Single-cell profiling of the developing mouse brain and spinal cord with split-pool barcoding. Science 360(6385), pp.176-182 (2018). 10.1126/science.aam8999

14. Srivatsan, S.R., McFaline-Figueroa, J.L., Ramani, V., Saunders, L., Cao, J., Packer, J., Pliner, H.A., Jackson, D.L., Daza, R.M., Christiansen, L. and Zhang, F. Massively multiplex chemical transcriptomics at single-cell resolution. Science 367(6473), pp.45-51 (2020). 10.1126/science.aax6234

15. Credle, J.J., Itoh, C.Y., Yuan, T., Sharma, R., Scott, E.R., Workman, R.E., Fan, Y., Housseau, F., Llosa, N.J., Bell, W.R. and Miller, H. Multiplexed analysis of fixed tissue RNA using Ligation in situ Hybridization. Nucleic acids research 45(14), pp.e128–e128 (2017). 10.1093/nar/gkx471

16. Schouten, J. et al. Relative quantification of 40 nucleic acid sequences by multiplex ligation-dependent probe amplification. Nucleic Acids Research 30(12) pp.e57 (2002). 10.1093/nar/gnf056

17. Rouhanifard, S. et al. ClampFISH detects individual nucleic acid molecules using click chemistry–based amplification. Nat Biotechnol 37, 84–89 (2019). 10.1038/nbt.4286

18. Pardo-Palacios, F.J., Wang, D., Reese, F. et al. Systematic assessment of long-read RNA-seq methods for transcript identification and quantification. Nat Methods 21, 1349–1363 (2024). 10.1038/s41592-024-02298-3

19. Lawlor, N. et al. Single Cell Analysis of Blood Mononuclear Cells Stimulated Through Either LPS or Anti-CD3 and Anti-CD28. Front Immunol. 12 636720 (2021) 10.3389/fimmu.2021.636720

20. Rade, M. et al. A time-resolved meta-analysis of consensus gene expression profiles during human T-cell activation. Genome Biology 24,287 (2023). 10.1186/s13059-023-03120-7

21. Domeier, P. et al. IFN-γ receptor and STAT1 signaling in B cells are central to spontaneous germinal center formation and autoimmunity. J Exp Med 213, 5 pp.715–732 (2016) 10.1084/jem.20151722

22. Elgueta, R. et al. Molecular mechanism and function of CD40/CD40L engagement in the immune system. Immunological Reviews 229 pp.152–172 (2009) 10.1111/j.1600-065X.2009.00782.x

23. Feng, H. et al. Interferon regulatory factor 1 (IRF1) and anti-pathogen innate immune responses. PLoS Pathog 17(1) pp.e1009220 (2021) 10.1371/journal.ppat.1009220

24. Sedger, L., McDermott, M. TNF and TNF-receptors: From mediators of cell death and inflammation to therapeutic giants – past, present and future. Cytokine & Growth Factor Reviews 25 (4) pp.453–472 (2014) 10.1016/j.cytogfr.2014.07.016

25. Szabo, P.A., Levitin, H.M., Miron, M. et al. Single-cell transcriptomics of human T cells reveals tissue and activation signatures in health and disease. Nat Commun 10, 4706 (2019). 10.1038/s41467-019-12464-3

26. González-Amaro, R. et al. Is CD69 an effective brake to control inflammatory diseases? Trends in Molecular Medicine 19(10) pp.625–632 (2013) 10.1016/j.molmed.2013.07.006

27. Holling, T. et al. Function and regulation of MHC class II molecules in T-lymphocytes: of mice and men. Human Immunology 65(4) pp.282–290 (2004) 10.1016/j.humimm.2004.01.005

28. Berkovits, B.D. and Mayr, C. Alternative 3′ UTRs act as scaffolds to regulate membrane protein localization. Nature 522(7556), pp.363-367 (2015). 10.1038/nature14321

29. Lee, S.H. and Mayr, C. Gain of additional BIRC3 protein functions through 3-UTR-mediated protein complex formation. Molecular cell 74(4), pp.701–712 (2019).

30. Mayr, C. Regulation by 3′-untranslated regions. Annual review of genetics 51, pp.171–194 (2017). 10.1016/j.molcel.2019.03.006

31. Moffitt, J.R., Hao, J., Wang, G., Chen, K.H., Babcock, H.P. and Zhuang, X. High-throughput single-cell gene-expression profiling with multiplexed error-robust fluorescence in situ hybridization. Proceedings of the National Academy of Sciences 113(39), pp.11046–11051 (2016). 10.1073/pnas.1612826113

32. Patro, R. et al. Salmon provides fast and bias-aware quantification of transcript expression. Nat Methods 14, 417–419 (2017). 10.1038/nmeth.4197

33. Langmead, B. and Salzberg, S.L. Fast gapped-read alignment with Bowtie 2. Nature Methods 9, pp.357–359 (2012). 10.1038/nmeth.1923

34. Wolf, F.A. et al. SCANPY: large-scale single-cell gene expression data analysis. Genome Biology 19(15) (2018). 10.1186/s13059-017-1382-0

