## Supplemental Figures and Notes for "HybriSeq: Probe-based Device-free Single-cell RNA Profiling"

### SUPPLEMENTARY MATERIALS

#### Supplemental Notes

##### Note 1

##### Non-split probe vs. Split probe

Taking inspiration from smFISH and multiplexed RNA FISH approaches we initially sought to make bulk measurements of transcripts with FISH probes by directly quantifying all probes adhered to cells with the assumption that specific hybridization would be much greater than nonspecific adhesion (Supplementary Fig. 1a). We found that nonspecific adhesion of probes to fixed cells was the same as probes targeting a specific transcript (Supplementary Fig. 1b). To overcome background adhesion, we specifically cleaved RNA-DNA hybrids with RNase H to release hybridized probes into solution which could then be captured (Supplementary Fig. 1a). RNase H mediated release led to specific signal above random probe binding that correlated with expected expression values from RNA-Seq (Supplementary Fig. 1c). We screened hybridization conditions and probe length for an optimal set of parameters but were only able to achieve a signal to background ratio of <200 for high expression targets (Supplementary Fig. 1d). While RNase H mediated specificity has the potential to be viable in a single cell approach, a nonnegligible amount of the released probes would likely be lost in the library prep before amplification leading to decreased signal. For these reasons, we opted for an alternative approach to specificity.

Adapting concepts from previous methods to detect RNA in fixed tissue sections with sequencing, we introduced a ligation step after hybridization in which adjacent probes hybridized to the same transcript would be ligated (Supplementary Fig. 2a) and probes nonspecifically adhered would not. With this approach, we were able to achieve a bulk signal that was > 1000-fold higher than nonspecific probe binding/ligation for moderately expressed targets as well as able to saturate the signal, which was not possible with unligatable probes (Supplementary Fig. 2b-c, Supplementary Fig. 1e). This contrasts with smFISH approaches which achieve specificity by the local enrichment of fluorescent signal by many labeled probes binding to transcripts of interest. We investigated multiple ligation conditions in bulk experiments and found that high ligase concentrations had no observable effect on the nonspecific probe ligation but did improve the accuracy of expression measurements (Supplementary Fig. 2d-e).

##### Note 2

##### Cell barcoding

To uniquely label transcriptomes of single cells with a barcode, we adapted steps from a previously demonstrated split and pool procedure with modifications. Initially, we tested barcode ligation conditions from previous reports in bulk but found that < 75% of our probes were ligated (Supplementary Fig. 3a). Because inefficiencies in each barcode ligation step contribute multiplicatively to decreasing the final detection efficiency as well as increased chance of barcode hopping, we screened a variety of barcode ligation conditions in bulk. A range of barcode/linker oligo concentrations and ligation times were assessed. Ligation for over 240 min at a barcode/linker concentration of over 200 nM did not significantly increase the barcode ligation for a single probe (Supplementary Fig. 3b). In addition to optimizing the ligation of barcodes, we also assessed the blocking step after each barcode ligation which blocks linker oligos from participating in ligation reactions during pooling (Supplementary Fig. 3c). We observed that blocking of the linker oligo with twice the barcode oligos concentration resulted in ~25% of free ends being blocked from ligation in the next round (Supplementary Fig. 3e). Suspecting that this might lead to excessive barcode hopping we changed our approach to quenching free ligateable ends with excess 3' oligo ends (Supplementary Fig. 3d), EDTA quenching the T4 ligase before pooling, and washing away unligated barcode oligos before the next ligation. Quenching of the free barcode oligos alone resulted in > 90% of free ends being blocked from ligation (Supplementary Fig. 3e). Quenching of the T4 ligase with EDTA along with washing resulted in a miss ligation rate of < 15% of free barcode oligos (Supplementary Fig. 3f). Washing also has the benefit of decreasing the chance of barcode hopping during PCR because there is no pre-amplification pull down like in other split pool approaches.

#### Supplementary Figures

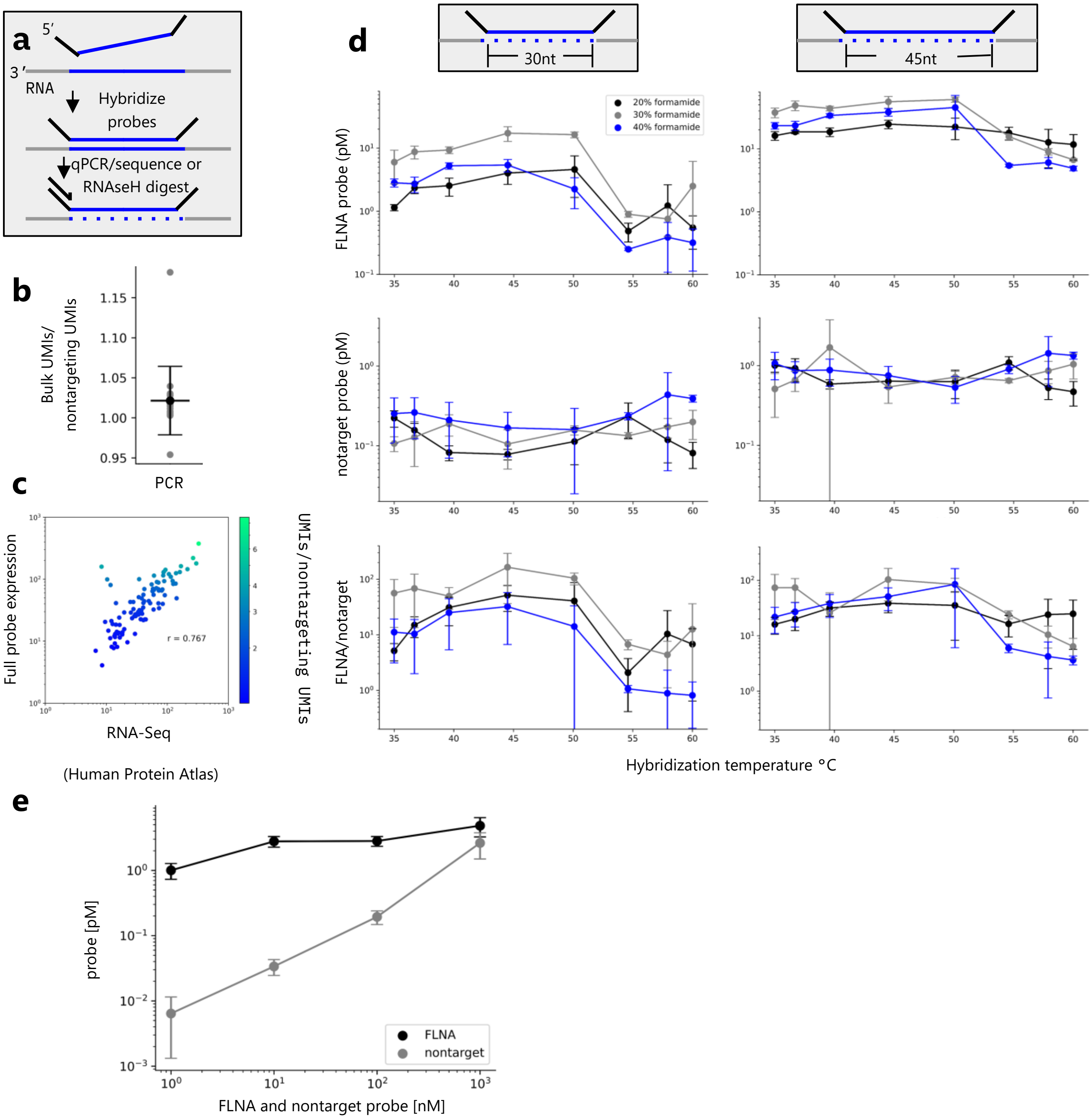

**Supplemental Fig. 1. Bulk Hybridization Conditions, Non-split Probes. a.** Schematic of the initial probe design and RNase H specificity method. Our initial approach was to hybridize non-split probes and qPCR/sequence them directly or release RNA-bound probes with RNase H. **b.** Bulk UMIs/nontarget probes(30nt) for probes targeting transduced fluorescent protein transcripts in HEK293 cells. Cells were directly loaded into a limited cycle PCR to amplify cell-associated probes. Probes were sequenced. Each point represents a unique probe. **c.** Scatter plot of bulk scaled expression values and RNA-Seq scaled expression values for genes measured with non-split probes (30 nt) released via RNase H. ~20 probes were used to detect the same transcript, and their UMIs were averaged together to yield an expression value. Each point represents a gene measured colored by average expression above nontargeting probes. Pearson correlation coefficient $r$ = 0.767 **d.** Hybridization condition optimization for non-split probes released via RNase H of lengths 30nt (left column) and 45nt (right column). Concentrations of RNase H-released probes targeting FLNA RNA or nontargeted in HeLa cells measured with qPCR. Error bars represent the standard deviation associated with technical replicates. % formamide is the concentration of formamide used in the hybridization buffer. **e.** The concentration of RNase H released probes (30 nt) targeting FLNA RNA or nontargeted in HeLa cells, measured with qPCR for 1-1000 nM probe hybridization concentration and 45 °C hybridization temperature.

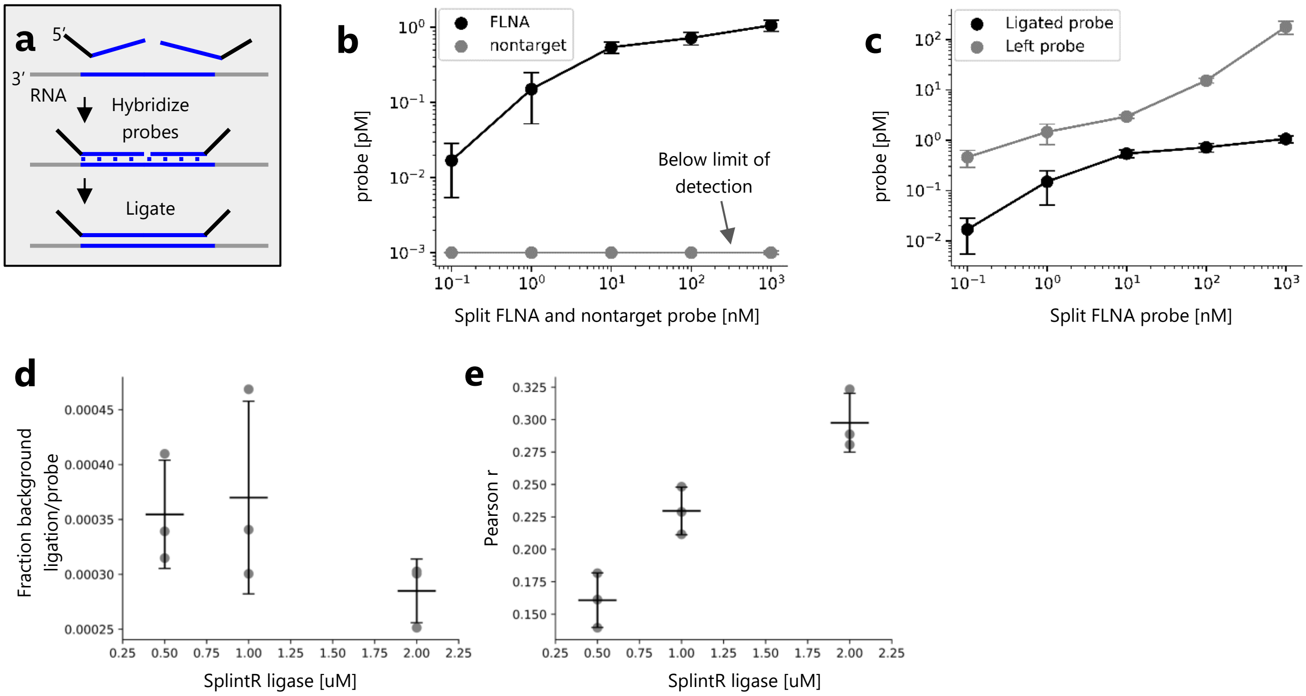


**Supplemental Fig. 2. Probe Ligation Optimization. a.** Schematic of split probe design and ligation specificity method. **b.** Bulk concentration of ligated probes targeting FLNA RNA or nontargeted in HeLa cells measured with qPCR for 0.1-1000 nM split probe hybridization concentration. For all hybridization concentration the nontargeting probes were below the limit of detection (0.001 pM). Error bars represent the standard deviation associated with technical replicates. **c.** Bulk concentration of ligated and left (unligated) probes targeting FLNA RNA in HeLa cells measured with qPCR for 0.1-1000nM split probe hybridization concentration. Error bars represent the standard deviation associated with technical replicates. **d.** Bulk fraction of misligation events/# of probes in the library. Misligation is the result of a left and right probe ligating that does not target adjacent regions of a transcript. Error bars represent the standard deviation associated with biological replicates n = 3. **e.** Pearson correlation coefficient (*r*) for correlation between bulk HybriSeq counts and RNA-Seq expression values for matched cell lines. Concentration refers to SplintR ligase used to ligate probes. Error bars represent the standard deviation associated with biological replicates n = 3.

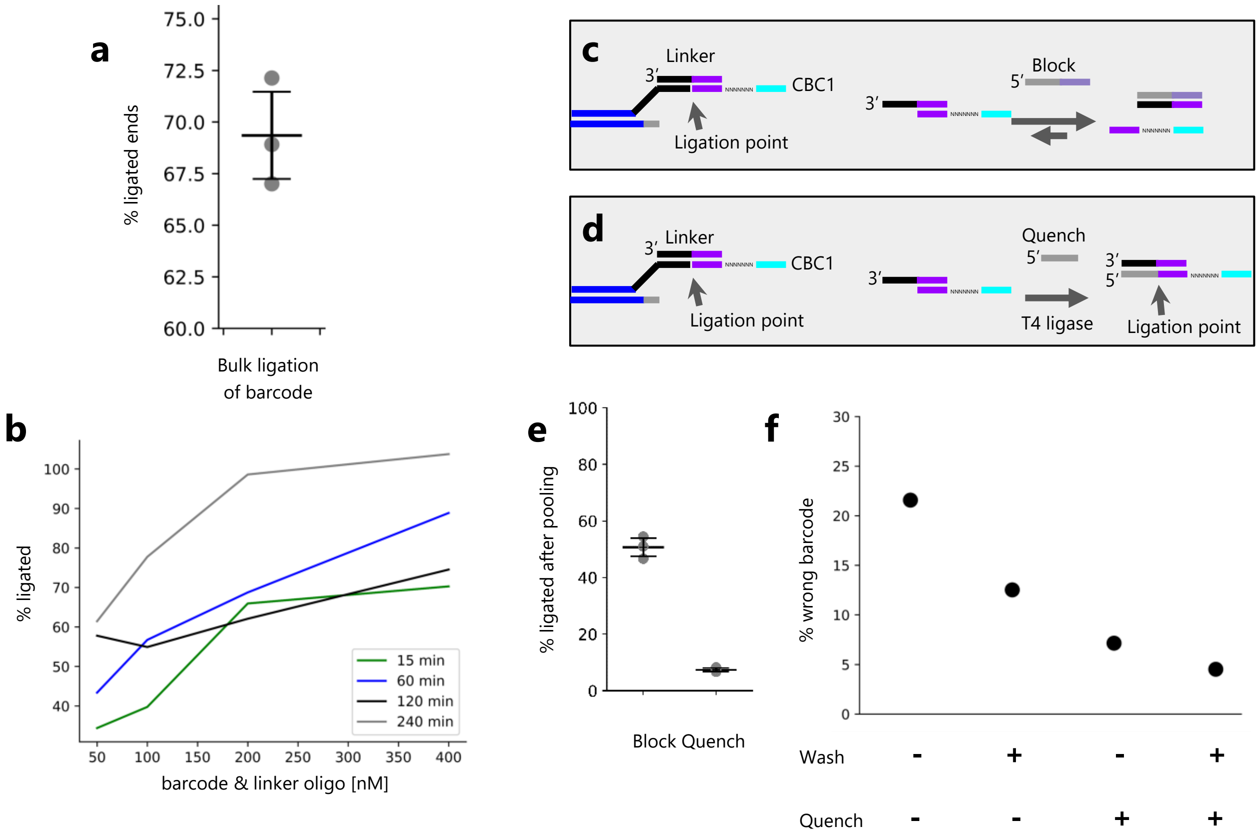


**Supplemental Fig. 3. Barcode Ligation Optimization. a.** Initial percentage of probe ligated ends for original round 1 barcode ligation conditions (15 minutes, 25°C, 200nM CBC & linker) measured in bulk with qPCR. Error bars represent the standard deviation associated with biological replicates n = 3. **b.** Percentage of probe ligated ends for round 1 barcode ligation for ligation times 15-240 min and barcode/linker concentration in reactions of 50 nM, 100 nM, 200 nM, and, 400nM. Measured in bulk with qPCR. **c.** Schematic of initial blocking strategy. Blocking oligos are used to displace linker oligos and prevent them from participating in ligation during subsequent rounds of cell barcoding. **d.** Schematic of HybriSeq quenching strategy. Quench oligos are used to ligate onto the ends of free unligated barcodes thereby preventing them from participating in subsequent ligation reactions. **e.** Barcode oligos/linkers were either blocked as in **c** or quenched as in **d**, cells with probes with free unligated 3’ ends were added and allowed to react with blocked or quenched barcodes for 2 hours and the fraction of ligated and unligated probe 3’ ends were measured with qPCR. Error bars represent the standard deviation associated with technical replicates. **f.** Cells went through 2 rounds of ligation-based barcoding with one of two barcodes in each round. In each pooling step, the unreacted or quenched other barcodes were added and allowed to react. Cells were either washed 2 times in EDAT solution after pooling or not washed and put into the next round of barcoding. The quantity of each possible barcode was measured with qPCR and the percent incorrect barcode was calculated for each condition.

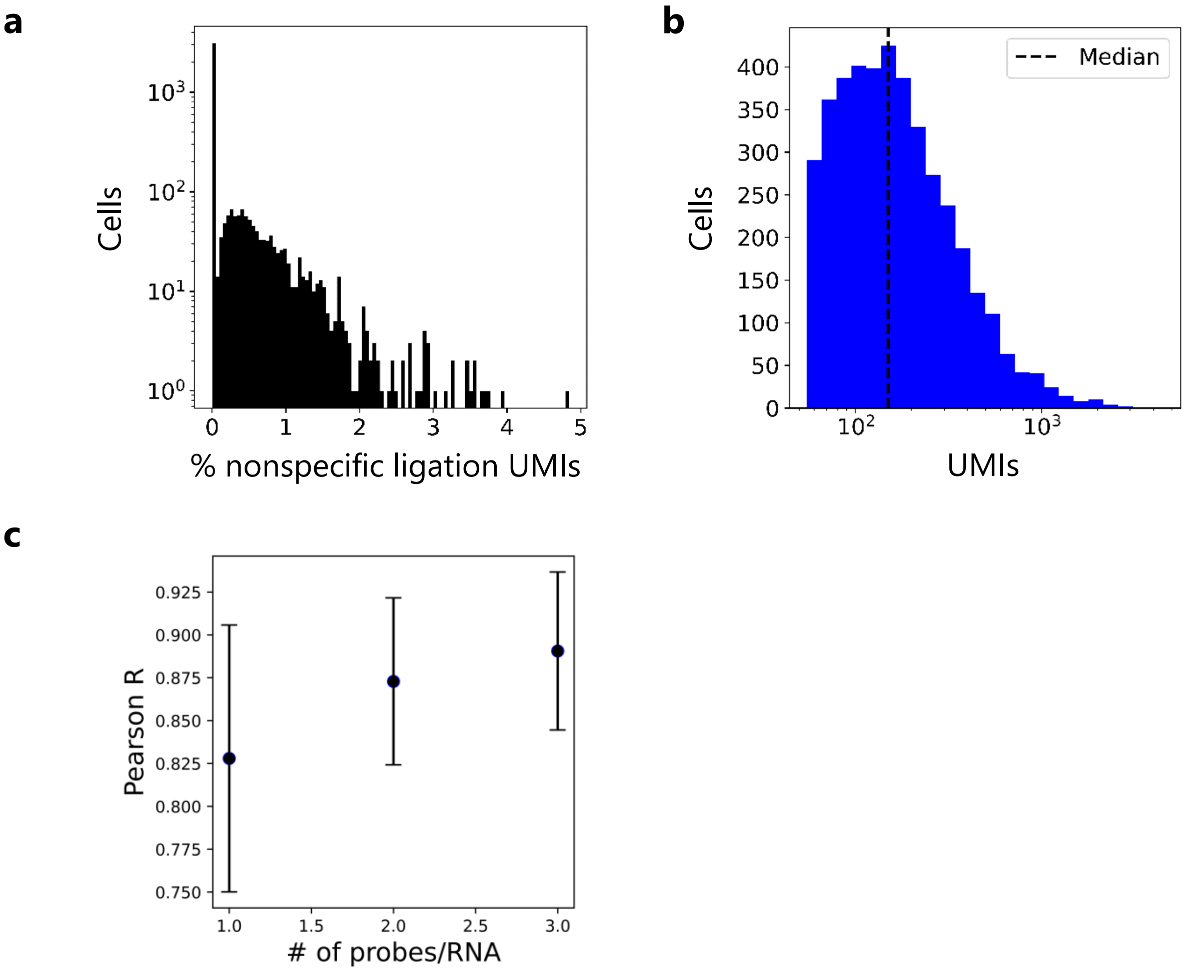


**Supplemental Fig. 4. Single-cell Validation. a.** Histogram of the percent of nonspecific ligation UMIs in cell mixing data. The average percent nonspecific ligation UMIs per cell is 0.20%. 74% of cells have 0 nonspecific ligation UMIs. **b.** Histogram of UMIs in cell mixing data. The median number of UMIs per cell was 150. **c.** Pearson correlation coefficients between bulk HybriSeq expression values and cell line matched RNA-Seq expression (TPM) for subsampled number of probes/gene used to make HybriSeq measurements. Error bars represent the standard deviation associated with each measurement obtained by bootstrap via sampling the unique probes used to calculate the correlation coefficient.

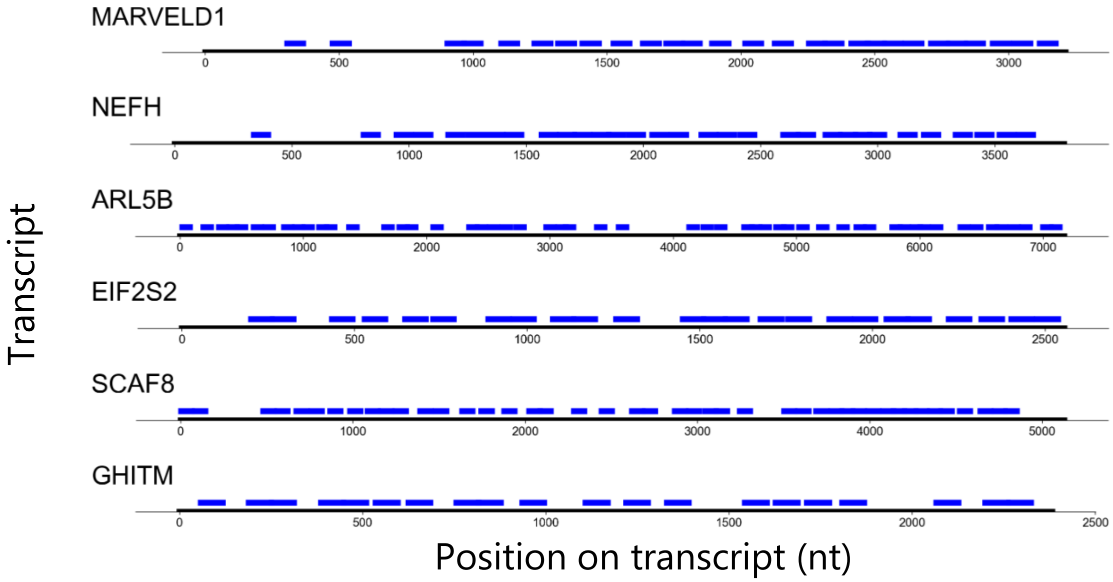


**Supplemental Fig. 5. Tiling Probes.** Tiling probes plotted on each transcript. ~50% of the transcripts are occupied by a probe.

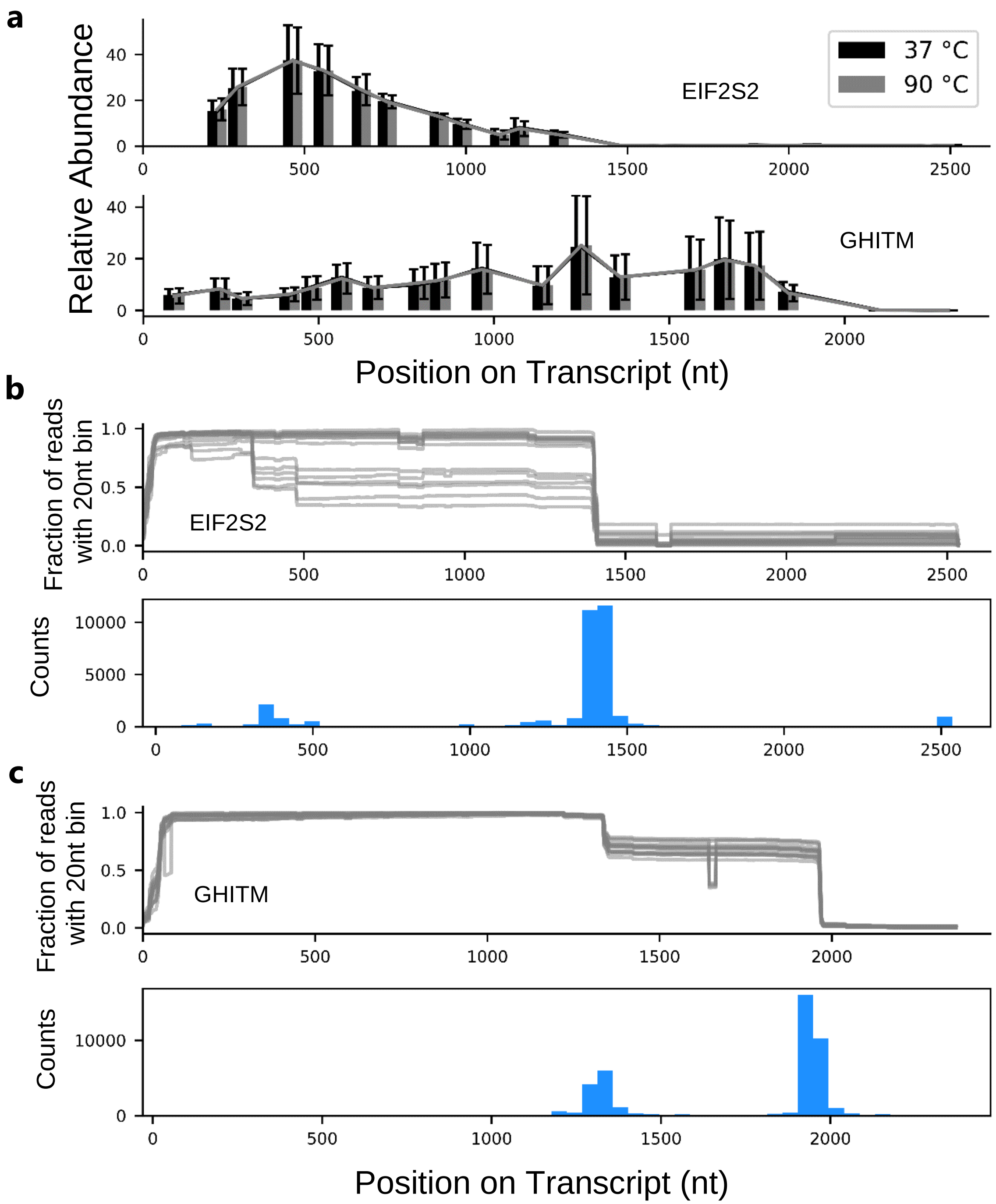


**Supplemental Fig. 6. *In vitro* probing of transcripts and Long-read RNA-Seq Genome Annotation Assessment Project (LRGASP) data. a.** Relative probe abundance for probes targeting EIF2S2 (top) and GHITM (bottom) in vitro for normal hybridization conditions (black) or denaturing conditions (gray) with standard deviation across three independent experiments. **b.** Top: fraction of EIF2S2 reads from 15 datasets which contain the 20nt segment of the annotated transcript from Ensembl annotated transcript ENST00000374980.3. Each line represents one of the 15 datasets. Bottom: histogram of read length from data above. **c.** Top: same as **a** for GHITM annotated transcript from Ensembl ENST00000372134.6. Bottom: same as **b** for GHITM.

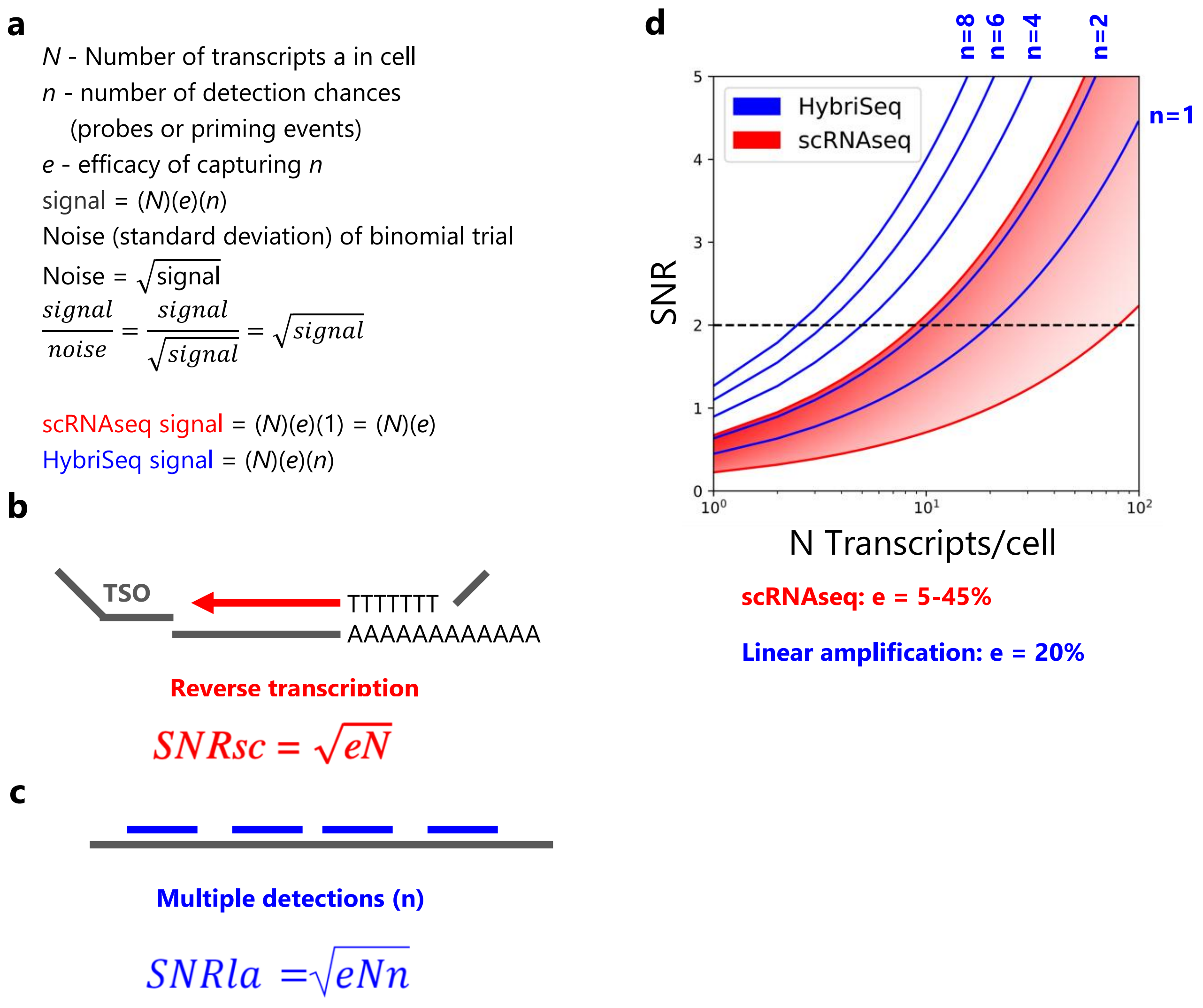


**Supplemental Fig. 7. SNR Model for HybriSeq with Multiple Probes. a.** Model of SNR in HybriSeq and scRNAseq measurements. The sampling of a specific transcript is assumed to be Poisson in nature. **b.** Poly(dT) priming on a polyadenylated transcript and template switching with template switching oligo (TSO). The theoretical SNR of the measurement is proportional to the square root of the efficiency of capturing *N* number of transcripts in a cell. **c.** Multiple detection possibilities with multiple probes/transcripts. The theoretical SNR of the measurement in HybriSeq is proportional to the square root of the efficiency of capturing n number of detection possibilities for *N* transcripts in a cell. **d** Theoretical measurement SNR for scRNAseq measurement made like in **b** with an efficiency of 5-45% and multiple detections as in **c** with an efficiency of 20%. With multiple detection possibilities or probes the minimum number of transcripts that can be detected with an SNR > 2 increases.

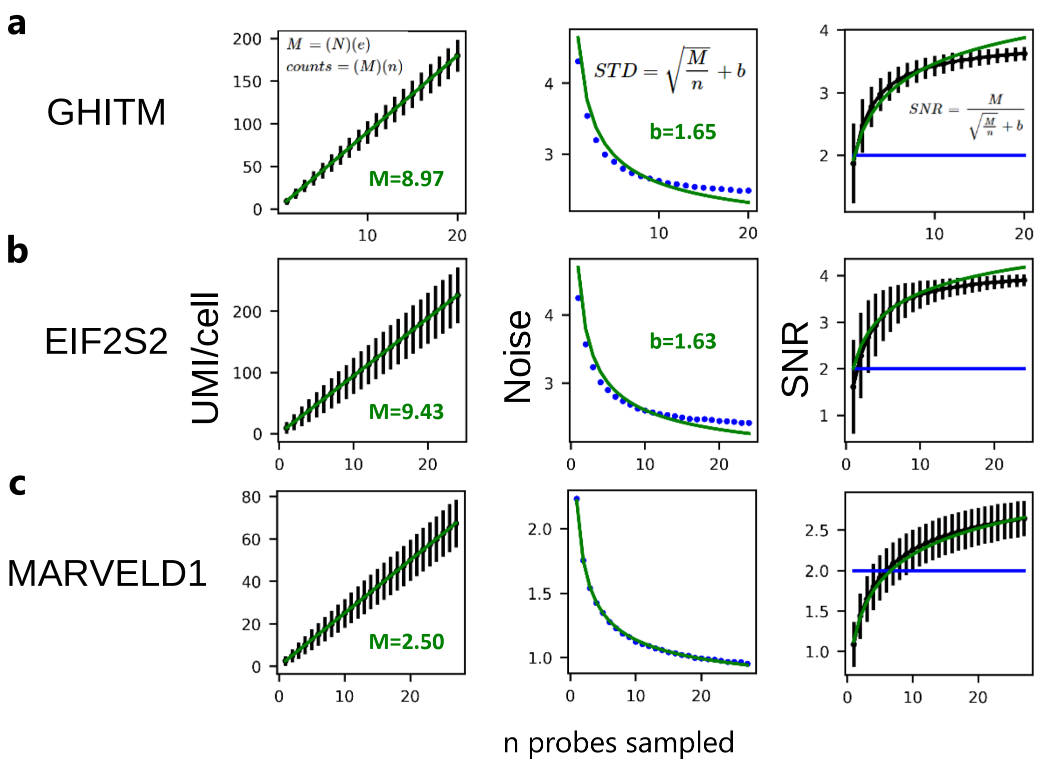


**Supplemental Fig. 8. Linear amplification with HybriSeq. a.** Left: Experimentally obtained values for average GHITM UMIs/cell for the sampled number of probes n. Error bars represent the standard deviation associated with each measurement obtained by bootstrap via sampling the unique probes used to calculate average UMIs/cell. The green line represents the fitted linear curve with slope *M*. Middle: Experimentally obtained values for the noise in expression measured with multiple probes *n*. The green line represents the fitted model from (Supplemental Fig. 7A) with the addition of a baseline variance to account for non-measurement-associated heterogeneity. Right: Experimentally obtained values for the measurement SNR measured with multiple probes n. Error bars represent the standard deviation associated with each measurement obtained by bootstrap via sampling the unique probes used to calculate average SNR. The green line represents the fitted model from the middle, and the blue horizontal line denotes SNR = 2. **b.** Same as **a** for EIF2S2. **c.** Same as **a** for MARVELD1.

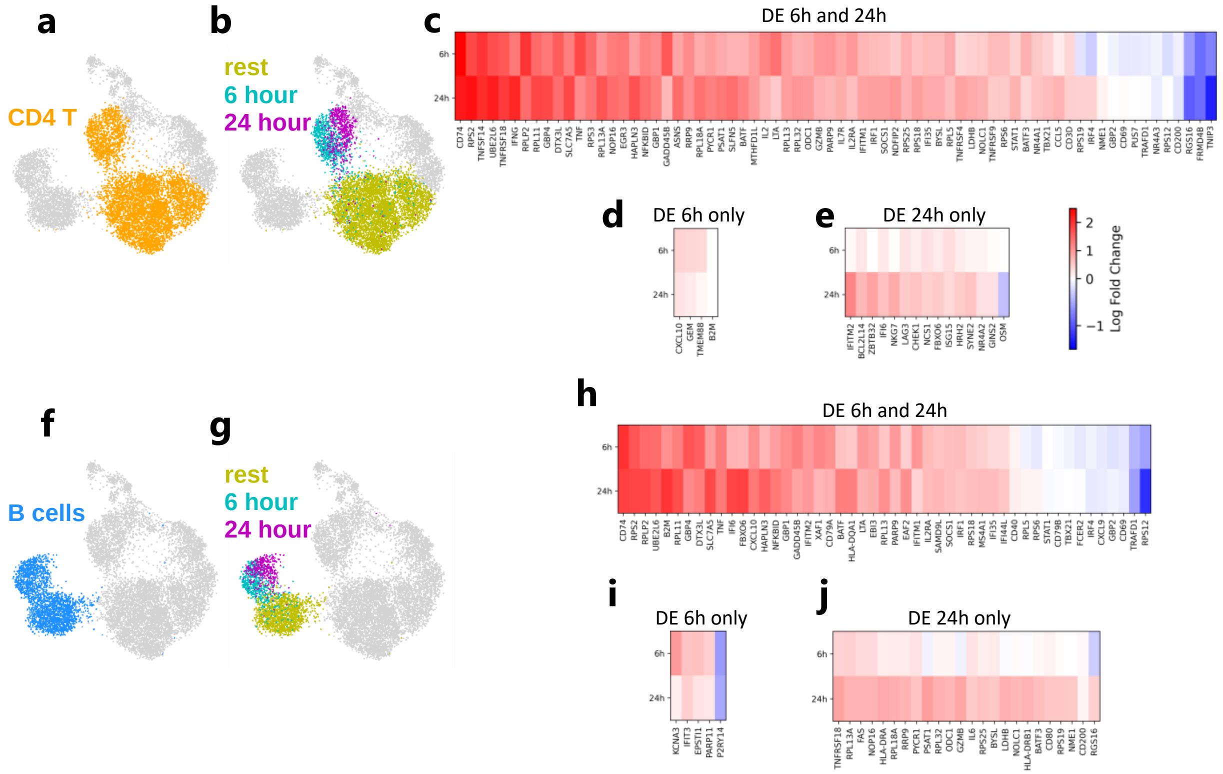


**Supplemental Fig. 9. Differentially expressed genes in CD4 T cells and B cells. a.** UMAP of PBMCs, CD4 T cells highlighted. **b.** UMAP of highlighted cells in **a** colored by stimulation condition. **c.** genes differentially expressed in CD4 T cells at 6 hours and 24 hours compared to unstimulated CD4 T cells (rest). **d.** genes differentially expressed in CD4 T cells at 6 hours only compared to unstimulated CD4 T cells (rest). **e.** genes differentially expressed in CD4 T cells at 24 hours only compared to unstimulated CD4 T cells (rest). **f.** UMAP of PBMCs, B cells highlighted. **g.** UMAP of highlighted cells in **f** colored by stimulation condition. **h.** genes differentially expressed in B cells at 6 hours and 24 hours compared to unstimulated B cells (rest). **i.** genes differentially expressed in B cells at 6 hours only compared to unstimulated B cells (rest). **j.** genes differentially expressed in B cells at 24 hours only compared to unstimulated B cells (rest).
